# Lifelong tissue memory relies on spatially organised dedicated progenitors located distally from the injury

**DOI:** 10.1101/2023.02.02.526841

**Authors:** Chiara Levra Levron, Mika Watanabe, Valentina Proserpio, Gabriele Piacenti, Andrea Lauria, Stefan Kaltenbach, Takuma Nohara, Francesca Anselmi, Carlotta Duval, Daniela Donna, Denis Baev, Ken Natsuga, Tzachi Hagai, Salvatore Oliviero, Giacomo Donati

## Abstract

It is believed epithelial cells that have participated in a wound repair elicit a more efficient but locally restricted response to future injuries. However here we show that the cell adaptation resulting from a localised tissue damage has a wide spatial impact at a scale not previously noticed. We demonstrate that away from injured site, after a first injury a specific epithelial stem cell population gives rise to long term wound-memory progenitors residing in their own niche of origin. Notably these progenitors have not taken part in the first wound healing but become pre-activated through *priming*. This adaptation differs from classical features of *trained immunity* previously shown to be adopted by other epithelial stem cells. Our newly identified wound-distal memory cells display a cell-autonomous transcriptional pre-activated state leading to an enhanced wound repair ability that can be partially recapitulated through epigenetic perturbation even in absence of an injury. Importantly, the harmful consequences of wound repair, such as exacerbated tumorigenesis, occur within these primed cells and follow their spatial distribution. Overall, we show that sub-organ scale adaptation of an injury relies on spatially organised and memory-dedicated progenitors, characterised by an epigenetic actionable cell state, that predisposes to tumour onset.

## Introduction

Forming the outer layer of organs, epithelia have predominantly a barrier function and are able to sense and adapt to environmental changes. The homeostatic integrity of these tissues is maintained through a continuous turnover ensured by stem cells (SCs), that can be compartmentalised in specific epithelial niches in skin^1–3^ as well as in those so-called transition zones, present in the epithelia of oesophagus, eye, anus, lung, stomach, and cervix ^4,5^. Upon tissue damage, however, the defined niche boundaries can be lost and each epithelial lineage resident nearby the injury, acquires cell plasticity that allows it to migrate towards the wound site, thus contributing to the re-epithelialization^6–11^.

In the last 5 years it emerged that epithelial cells adapt to a local stressful event, such as wound, though the establishment of a chromatin memory to respond faster to an eventual similar challenge ^12,13^. Nevertheless, the lineage specificity of wound memories, through a direct comparison of different epidermal cell populations, has not been dissected yet^14^.

The pioneer work by the Fuchs laboratory showed that the stem cells located in close proximity of the injured tissue can be trained, while participating in the repair, suggesting an extremely locally restricted potential of wound memory^12,13^. Moreover, beside the positive effect of memory on regenerative potential^15^, negative consequences such as cancer^16^ might be related to it. In this context, it would be of translational interest to understand the spatial distribution and extent of wound memory in epithelial cells located distally from the repaired area and to characterise the full impact on the epithelium, long term.

It has been demonstrated that, in parallel to immune system, epithelial cells too exhibit *trained wound-memory* of an injury. After a wound event, these cells on the damaged area can establish chromatin memory that, following re-set of homeostasis, is kept transcriptionally dormant. However, this chromatin state allows a quick reactivation in the event of an eventual lesion^12,13,15^. Importantly, innate immune adaptation has been categorised to distinguish distinct processes including, the so-called *trained immunity* and *priming*^17^. Differently from *trained immunity* where the activation status goes from active to inactive following resolution, *priming* connotes an activation state that never turns off even when the stimulus ceases. Currently, it is unknown if other adaptation programs, such as priming, are opted by epithelial cells^14^.

Here, in the context of two consecutive skin injuries, we combined lineage tracing with single cell analysis to comprehensively understand the spatial extent of wound memory and the full spectrum of the adaptive responses of epithelial cells (i.e. *trained wound-memory* vs *wound priming*) within the same lineage and amongst different lineages. We show that specific SCs give rise to wound-primed progenitors that exist in a wide undamaged area distally from the damaged zone while within their original epidermal niche. Mechanistically, we demonstrate that transcriptional de-repression is functional for memory onset. Finally, the primed memory that is established outside the wound site has detrimental consequences on ageing-related diseases, such as skin cancer. Altogether, our unexpected results drastically change the assumption relative to the spatial distribution of memory cells by highlighting the existence of memory progenitors located far from the injury, in undamaged areas. The adaptation program co-opted by wound-distal memory cells represents the first description of priming in epithelial context.

## Results

### Wound healing educates Lrig1 stem cell progeny distally from the injury in undamaged areas

Hair follicle (HF) cells are recruited during wound healing and the actual contribution to epidermal repair is specific for each cell lineage^9^. Recently, the existence of an epigenetic wound memory has been assessed for hair follicle stem cells (HFSCs)^15^, but the precise lineage identity, the spatial distribution and the spectrum of adaptation programs acquired by the memory cells are still unknown (Fig.1a). We focused on three well-characterised cell populations compartmentalised in HF: Lrig1^+^ SCs mainly localise in the HF junctional zone (JZ) and are involved in the maintenance of the sebocytes, sebaceous ducts and the infundibulum (INFU); Gata6^+^ are committed-to-differentiation and differentiated duct cells of the upper pilosebaceous unit; Lgr5^+^ SCs are confined in the lower HF and are responsible for its cyclical growth^11,18,19^.

**Fig.1:**
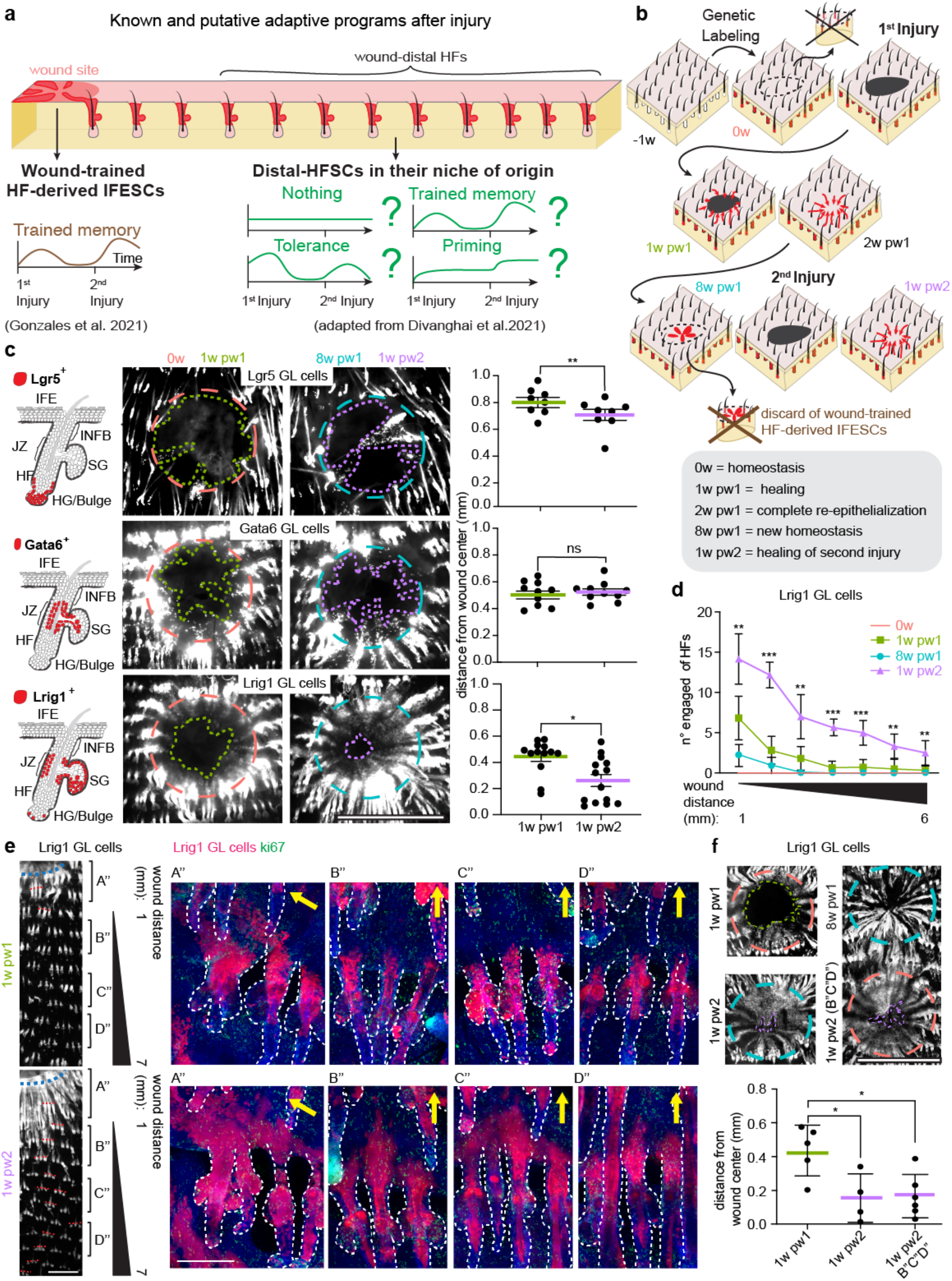
Wound memory in hair follicle lineages and its spatial extension. **(a)** Schematic representation of the state of art and the study’s objectives. On the left the repairing hair follicle (HFs) in proximity of the wound and the HF-derived newly formed epidermis, focus of Gonzales et al. study ^15^, are reported. On the right distal HFs are made up of HF cells in their niche of origin. Their adaptation program to wound will be the focus of this study. **(b)** Two consecutive skin injuries model: (−1w) genetical labelling (GL) through tamoxifen application; (0w) homeostasis; first injury; (1w pw1) healing phase, one week post first wound; (2w pw1) complete re-epithelialization, two weeks post first wound; (8w pw1) new homeostasis, eight weeks post first wound, and second injury; (1w pw2) second healing phase, one week post second wound. **(c)** The localisations of Lrig1^+^, Lgr5^+^ and Gata6^+^ cell populations in the HF are shown on the left. Whole-mount of epidermis (red channel) showing GL tdTomato^+^ cells exiting from HFs (central panels) and quantification of the distance from wound centre (right panels) are reported for each population. Dashed circles indicate the original wound perimeter (orange for 0w and light blue for 8w pw1); dashed lines (green for 1w pw1 and purple for 1w pw2) underline the migratory front of GL cells. n=8-14 wounds. Mean ± SEM is plotted. **(d, e)** HF engagement, as the exit of Lrig1 GL cells from HFs into the interfollicular epidermis (IFE), is shown. Number of engaged HFs at different distances, up to 7 mm away from wound. n=6-9 wounds (d). Stereoscopic images (red channel) of engaged HFs at 1w pw1 and 1w pw2 (left) and confocal images (right) in different locations: A’’, B”, C” and D” as indicated. Yellow arrows the position of the wound site (e). **(f)** Upper panel: pictures of 1w pw2 healing of a second overlapping wound (1w pw2) (left) or an injury inferred in B”C”D” region (1w pw2 (B”C”D”)) (right). Dashed circles indicate the original wound perimeter at time 8w pw1 (light blue) or 0w (orange) while the lines underline the migration front of the labelled cells at 1w pw2 (purple) or 1w pw1 (green). Lower panel: distance from wound centre is quantified in the indicated settings. n=4-6 wounds. P-value: *** 0.0001 to 0.001, ** 0.001 to 0.01, * 0.01 to 0.05, ns ≥ 0.05. Mean ± SD is plotted if not differently indicated. Scale bars: 2 mm (c, f); 150 μm (**e**).

To understand if their progenies elicited a wound-induced memory, we genetically labelled (GL) Lrig1^+^, Lgr5^+^, and Gata6^+^ HF cells (Extended Data Fig.1a,b) in adult mice. We used a two consecutive injuries model to assess memory in tail skin (Fig.1b), where, in the absence of tissue contraction (Extended Data Fig.1c,d), the second injury heals faster than the first one (Extended Data Fig.1e,f). Briefly, at time 0w we performed a first full thickness wound and 8 weeks after (8w pw1), when a new homeostasis is set (Extended Data Fig.1g-i), we induced a second identical and overlapping injury (Fig.1b).

It has been shown that when HFSCs leave their niches and take up residence within the repaired tissue, they have enhanced repair ability in case of a second assault thanks to the establishment of epigenetic memory^15^. Importantly, in our experiments, the second wound perfectly overlaps the first one, allowing the removal of the HF-derived IFESCs (Fig.1b). These cells, activated during the first wound and residing in the newly formed epidermis, were the focus of Gonzales et al.^15^. Thanks to this strategy we investigated the memory of the HFSCs that remain localised in their original niche in the HF, without contributing to the repair of the IFE.

Despite the contribution of Lrig1 GL cells is quantitatively higher than Lgr5 GL cells, both progenies show enhanced re-epithelialization ability during the second healing (1w pw2) compared to the first one (1w pw1), while Gata6 GL cells do not (Fig.1c and Extended Data Fig.1j). These results indicate that differentiating HF cells (here represented by Gata6^+^) do not show wound adaptation while HFSCs, independently of their epidermal lineage, do.

The spatial extent of the wound memory is unknown, although it has been recently shown that the cell contribution to skin full thickness wound repair is spatially restricted to less than 1 mm away from the injury site^20^. Consistently with this observation, during the first healing, the wound-engaged HFs (defined as HF in which GL Tomato^+^ cells move from their homeostatic HF niche into IFE) are mainly localised in the close surroundings of the injury (Fig.1c and Extended Data Fig.1j). However, at 1w pw2 this phenotype exists at least 7 mm away from the injury site, exclusively for Lrig1 progeny (Fig.1d,e and Extended Data Fig.1k,l). This unexpected finding suggests that wound experience educates Lrig1 GL cells resident in distal HFs that did not directly contribute to the IFE repair. To confirm this, we performed a second injury distant from the site of the previously healed area (zones B”C”D”) in order to directly challenge distal HFs that haven’t been engaged during the first healing (Figure 1F). Strikingly, in this setting the second healing rate is enhanced, confirming that memory is acquired by follicular Lrig1 GL cells also at great distance from the first wound in undamaged areas.

Thus, we show that, as a consequence of a lesion, different SC lineages acquire a wound memory if located in close proximity of the injury. However, exclusively the Lrig1^+^ SC progeny is wound-educated within the HFs that localise distally from injury, up to ∼ 7 mm.

### Distal memory is cell-autonomous

Since only wound-distal Lrig1 GL cells, but not Lgr5 ones, show adaptive behaviour to injury, to decipher their transcriptional mechanisms we compared the expression profile of Lrgi1 and Lgr5 GL tdTomato^+^ sorted cells in a longitudinal population-specific RNA-Seq experiment. Consistently with their different adaptive phenotype, the analysis shows that the two HFSC lineages have specific wound-associated transcriptional programs (Extended Data Fig.2a). We defined as memory genes those genes that were deregulated during the first healing and whose deregulation was of greater magnitude after the second injury. Lrig1 GL cells have a higher abundance of memory genes (almost 50% of the total differentially expressed genes) respect to Lgr5 GL cells (Fig.2a). Gene Ontology (GO) analysis of them suggests “Cell polarity/Migration”, a major cell phenotype in wound healing^20,21^, as a feature of the wound-educated Lrig1 GL cells (Fig.2b). This corroborates previous histological data about the enhanced migration of Lrig1 GL cells during the second healing (Fig.1c), that is not robustly sustained by proliferation (Fig.2c).

**Fig.2:**
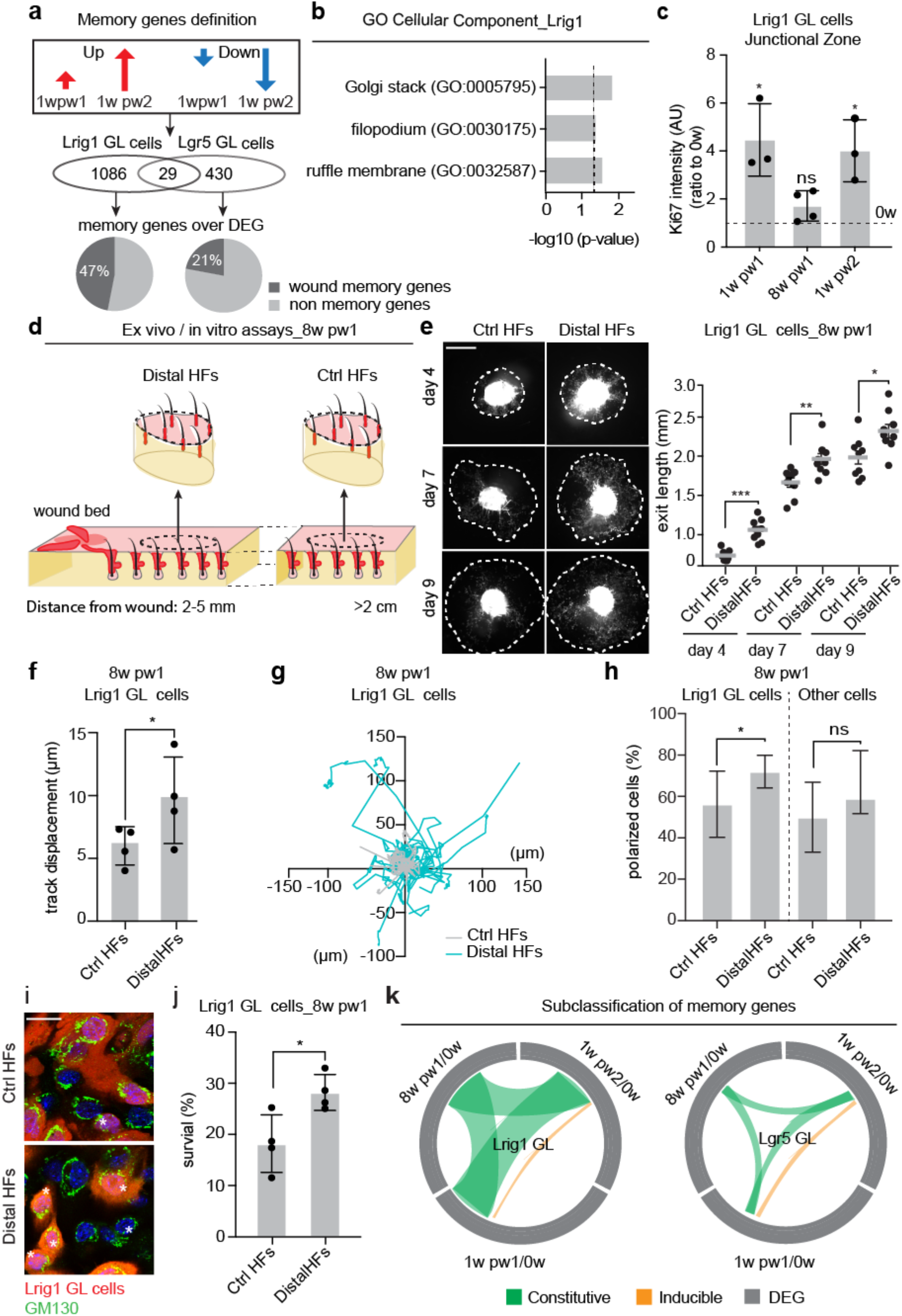
Cell-autonomous transcriptional program enhanced fitness of wound-experienced Lrig1 progeny. **(a)** Memory genes definition (logFC (1w pw2) > logFC (1w pw1)) and Venn diagram reporting the number and the percentage of inferred memory genes over the total differential expressed genes (DEG) from Lgr5 and Lrig1 GL cells. **(b)** Enriched GO terms for memory genes in Lrig1 GL cells. - Log10(p-value) is plotted. Dashed line indicates significance. **(c)** Quantification of the Ki67 index in junctional zone (JZ) of Lrig1 GL cells at indicated time points, normalized to 0w. n=3-4 wounds. **(d)** Ex vivo and in vitro experimental settings at 8w pw1: Lrig1 GL skin biopsies from wound-educated Distal HFs (within 5 mm from wound site) and from Ctrl HFs, outside from memory zone (distance greater than 2 cm) are compared. **(e)** Images of ex vivo migration of Lrig1 GL cells (left panel) and quantification of the migration (exit length) (right panel). Dashed lines mark the migration front of epidermal cells. Mean ± SEM are reported. Scale bar: 2 mm. n=9 skin explants. **(f, g)** Time lapse migration assay. Track displacement (μm) (f) and representative normalised start position graph (g) of Lrig1 GL cells movement over 16 hours of time lapse. n=4 mice. **(h, i)** Cis-Golgi (GM130) distribution in cells exiting from skin explant. Percentage of polarised cells (marked by asterisk) in GL tdTomato^+^ and tdTomato^-^ cells (h) and representative pictures (i) are shown. n=4 explants; Scale bar: 20 μm. **(j)** Surviving Lrig1 GL cells (Ctrl HFs or Distal HFs) as ratio between day 2 and day 0. n=4 mice. **(k)** Circular ideogram plot (CIRCOS) of shared DEG (grey) between 1w pw1, 8w pw1 and 1w pw2 (green) or between 1w pw1 and 1w pw2 only (yellow), in Lgr5 and Lrig1 GL skins. P-value: *** 0.0001 to 0.001, ** 0.001 to 0.01, * 0.01 to 0.05, ns ≥ 0.05. Mean ± SD is plotted if not differently indicated.

To dissect if distal memory is cell-autonomous, we collected skin biopsies at 8w pw1 from 2-5 mm far from the wound margins, where distal-educated Lrig1 GL cells reside (distal) or control zones (ctrl) (more than 2 cm far away from wound, where engaged HF are not detected) and performed ex vivo migration assay^13^ (Fig.2d and Extended Data Fig.2b,c). Lrig1 GL cells that experienced the wound display a higher in vitro migratory ability (Fig.2e and Extended Data Fig.2d,e), as also confirmed through time lapse imaging (Fig.2f,g and Extended Data Fig.2f) and the expression of the cell-polarity marker^22^ (Fig.2h), as well as adhesion and survival (Fig.2i and Extended Data Fig.2g). Taken together, these data indicate that Lrig1 GL cells resident distally from the lesion acquire enhanced repair capabilities that are intrinsic property of the cells.

Focusing on the new homeostatic state (8w pw1) in Lrig1 and Lgr5 GL epidermis, the memory genes display two distinct expression patterns (Fig.2j and Extended Data Fig.2h,i): (a) a constitutive pattern where the genes remain deregulated in between the two assaults; (b) an inducible trend where, after wound resolution, the original homeostatic expression is restored (0w). Interestingly, in Lrig1 GL cells almost 90% of the memory genes belong to the “a” type. Since 8w pw1 clearly represent a newly established homeostasis (Extended Data Fig.1), we hypothesised that the constitutive but intermediate (between the actively healing states) transcriptional activation of memory genes evidenced at 8w pw1 (Fig.2k and Extended Data Fig.2h,i) might be due to the existence of *wound priming* in a sub-population of Lrig1 SC progeny.

### A Lrig1 stem cell-derived population is wound-primed in distal HFs

To better dissect the transcriptional basis of wound memory taking in account the heterogeneity of cells in HF niches^23^ we performed single cell RNA-Sequencing (scRNA-Seq) by Smart-seq2 of ∼700 Lrig1 GL cells at four time points: homeostasis (0w), 1 week post first wound (1w pw1), new homeostasis (8w pw1) and 1 week post second wound (1w pw2) (Fig.3a and Extended Data Fig3a). Unsupervised clustering analysis (Fig.3b) and cluster marker identification (Extended Data Fig.3b) distinguish cells according to spatial location and differentiation stage in known epidermal niches as summarised in Fig.3c. Cell-cycle phases classification highlights the proliferative clusters (Fig.3d). The integration of our data with scRNA-Seq data from Joost et al.^23^, that provide markers for cells of epidermal niches and differentiation status, confirm our predictions on the epidermal compartments and differentiation direction of each cell cluster (Fig.3e). Pseudotime analysis on six trajectories as in Extended Data Fig.3c followed by GO analysis of the upregulated genes at the end of each trajectory shows enrichment of differentiation-related GO terms (Extended Data Fig.3d,e), corroborating the differentiation direction defined by the pseudotime (Fig.3c). Therefore, the data indicate that: (i) cluster 9 represents differentiated keratinocytes of the lower HF, (ii) cluster 13 is made by sebocytes, (iii) cluster 2 is formed by sebaceous and junctional zone duct cells, (iv) cluster 4 represents the INFU duct cells and (v) cluster 12 is composed by differentiated IFE cells. As previously suggested^18^, our scRNA-Seq data integrated with data from Dekoninck et al.^24^ confirm that Lrig1 GL cells can only contribute to the healing with only one of the two differentiated IFE lineages^25^, namely the interscale differentiation (Fig.3f and Extended Data Fig.3f).

**Fig.3:**
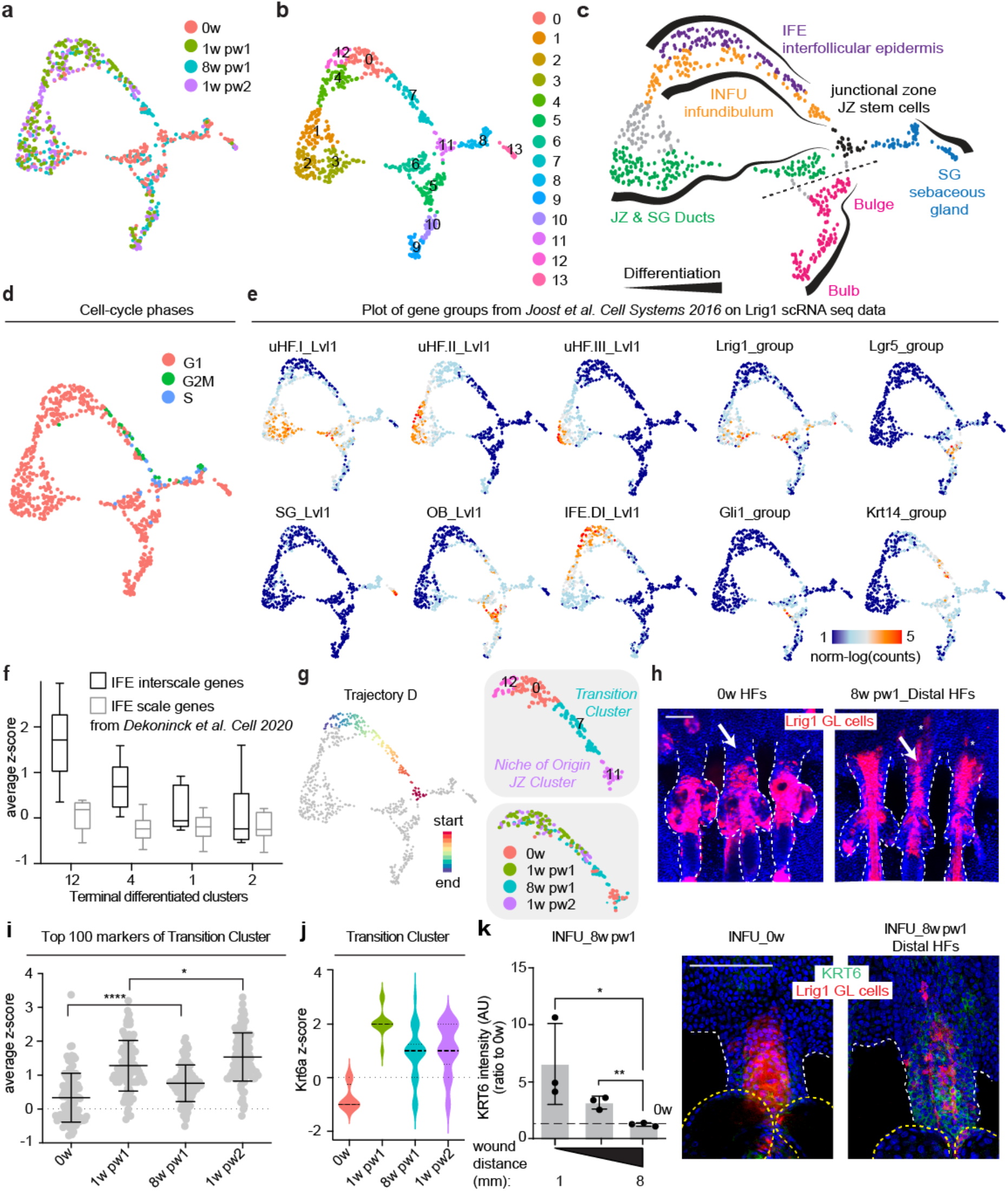
Identification of wound-primed cells in the infundibulum with a pre-activated transcriptional program. **(a)** UMAP of single cell data of Lrig1 GL cells from 0w (red), 1w pw1(green), 8w pw (light blue) and 1w pw2 (purple). **(b)** Unsupervised clustering of single cells data. **(c)** Summary illustrating epidermal compartments and differentiation in Lrig1 GL single cell data. Cells were coloured according to the epidermal compartment they belong to. Dashed line identifies the homeostatic lineage boundary between upper and lower HF (Page et al., 2013). **(d)** UMAP of cells coloured by assigned cell-cycle phase. (**e)** Plot of gene set expression from Joost et al. study^23^. **(f)** Whisker plot of average z-score for scale and interscale gene groups from Dekoninck et al. study^24^ in the epidermal terminal differentiation clusters of each trajectory. Median with 25^th^ and 75^th^ percentiles is plotted. **(g)** Trajectory D is coloured by time points and clusters. **(h)** Epidermal whole-mount images of Lrig1 GL tdTomato^+^ cells occupancy of the infundibulum (INFU) at 0w or in the distal HFs at 8w pw1 (at ∼ 5 mm from wound site). Asterisks mark differentiated IFE cells. **(i)** Plot of the average z-score of the top 100 markers of *Transition Cluster* (cluster 7) showing the intermediate transcriptional state at 8w pw1. **(j)** Violin plot of Krt6a expression in cells from *Transition Cluster* in each time point. **(k)** Quantification of Krt6 levels in the INFU at 8w pw1, at different distances from the wound site (ratio to 0w is plotted) (left) and whole-mount staining pictures at 0w and 8w pw1 (right) are shown. n=3 mice. P-value: **** < 0.0001, ** 0.001 to 0.01, * 0.01 to 0.05. Mean with SD is plotted if not differently indicated. Scale bars: 100 µm (h, k).

Trajectory D is the most interesting in terms of wound-induced plasticity. Indeed, Lrig1^+^ SCs acquire plasticity while leaving the *Niche of Origin-JZ Cluster* (cluster 11) and move along the trajectory in the *Transition Cluster* (cluster 7) towards differentiated IFE (cluster 0), where they contribute to the repair (Fig.3g). *Transition Cluster* contain cells that reside in the INFU at the starting point of the trajectory and then move towards IFE at the end of it, as suggested by the expression of the INFU marker Postn^23^ (Extended Data Fig.3g). Comparing the two homeostatic stages (0w and 8w pw1), we notice an unexpected increased in the number of Lrig1 GL cells in the INFU at 8w pw1, the newly established homeostasis 8 weeks after wound experience (Fig.3g,h). Since the infundibular sub-cluster expresses at 8w pw1 intermediate levels of the cluster markers genes in-between homeostatic state 0w and healing conditions (1w pw1 and 1w pw2) and the second induction at 1w pw2 reached greater transcriptional levels in comparison to 1w pw1 (Fig.3i), we deduced the existence of *priming* adaptive program, as described for immune cells^17^. This is also confirmed by GSEA ranking of “Cell activation” signature (Extended Data Fig.3i) and indeed Krt6 as a marker of wound-activated epidermal cells^26^ resulted as one of the primed transcript (Fig.3j and Extended Data Fig.3h).

Since our previous histological data indicated the Lrig1^+^ SC progeny can be educated to react faster to a second wound, even if distally located from the first healed injury (Fig.1d,e), we analysed the spatial expression pattern of Krt6. The histological validation of Krt6 confirms the spatial gradient of memory in the Lrig1 GL cells resident in INFU (Fig.3k), highlighting that the wound primed cells also exist distally from the injury.

Overall, these results indicate that, consequently to an injury, Lrig1^+^ SC-derived cells occupy the INFU where they remain transcriptionally pre-activated even when the damage has been resolved and a new homeostasis has been re-set. We show for the first-time the existence of *wound priming* as a memory of tissue repair in epithelial context, in sites a significant distance from the wound.

### Characterisation of distal priming

To dissect the transcriptional basis of *wound priming* in the second homeostasis (8w pw1), we characterised the expression profile of the cells in trajectory D (cluster 11, 7, 0 and 12), separating the first healing process (0w and 1w pw1) from the second one (8w pw1 and 1w pw2). The heatmap highlights the existence of cell plasticity genes that are specifically induced at higher levels during the second healing in the *Transition Cluster* (Fig.4a,b). The GO analysis detects enrichment for “Actin cytoskeleton”, “Epithelial-to-Mesenchymal Transition” (EMT) and “Focal adhesion” and is consistent with the cell-autonomous enhanced migratory abilities previously described (Fig.2 and Extended Data Fig.2), as well as Glycolysis and mTORC1 Signalling related terms (Fig.4b (right)).

**Fig.4:**
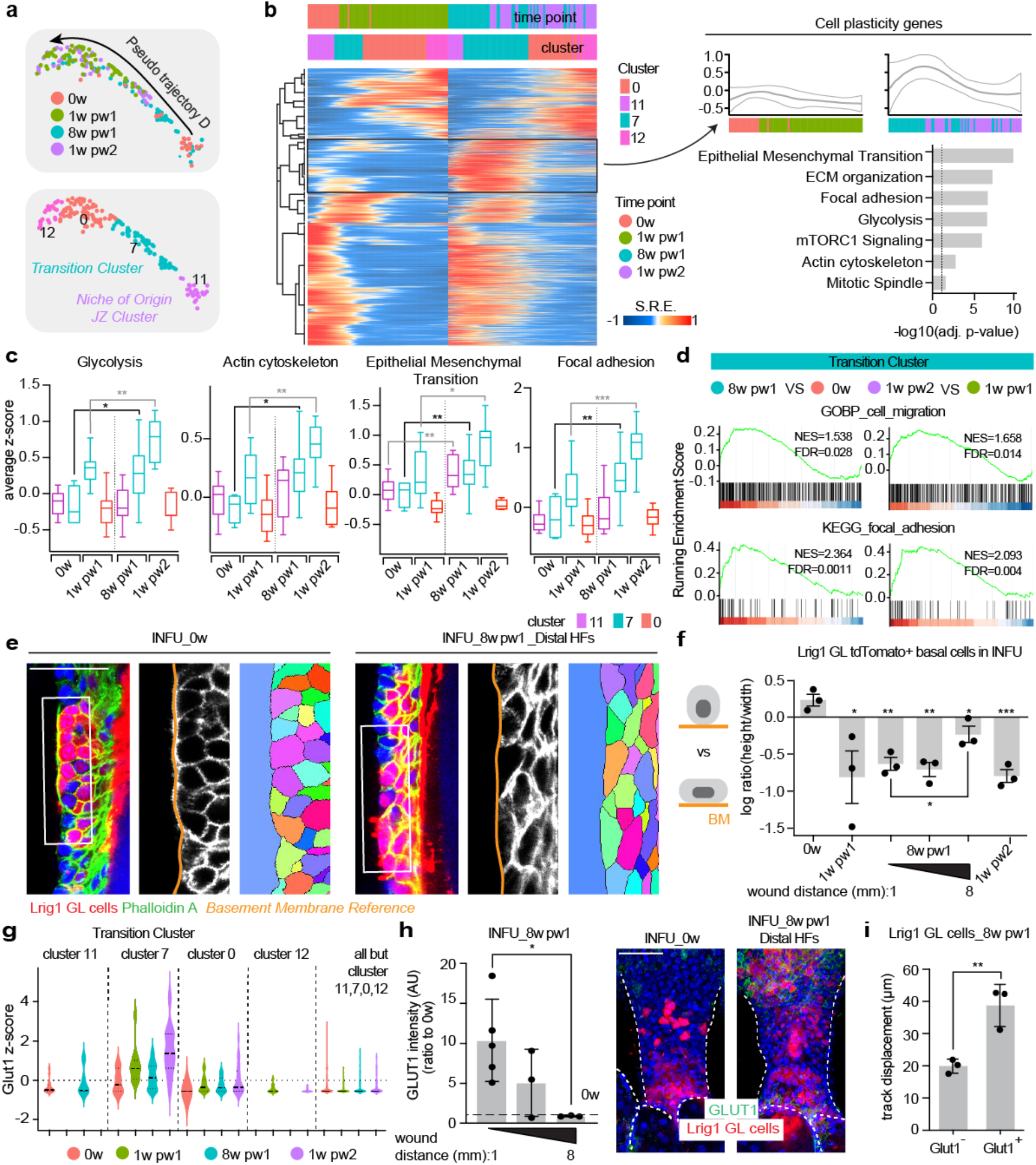
Spatial distribution and cell state characterisation of wound-primed cells. **(a)** UMAP of trajectory D with time point (up) and clusters (down). **(b)** Comparison of first and second healing. Left panel: Smoothed Relative Expression (SRE) of deregulated genes in trajectory D. Clusters and time points are indicated above and transiently induced cell plasticity genes are highlighted. Right panel: the average z-score expression of cell plasticity genes (up) and GO analysis (down) are shown. -log10 of the adj. p-value is plotted. The dashed line underlines significance. **(c)** Cells in trajectory D are divided for time points and clusters and the average z-score of the indicated GO term is plotted in the whisker plot. Median with 25^th^ and 75^th^ percentiles is plotted. **(d)** GSEA ranking of the indicated gene signature using differential gene expression from 8w pw1 vs is 0w (left panel) and 1w pw2 vs 1w pw1 (right panel) comparisons in cluster 7. **(e, f)** Single stack of epidermal whole-mount stained with phalloidin A (green in left panel), relative cell segmentation (right panel) (e) and quantification of cell shape as the ratio between height and width (f) in Lrig1 GL tdTomato^+^ cells in the infundibulum (INFU) at 0w and 8w pw1 (at ∼ 5 mm from wound site). n=3 mice. **(g)** Violin plot of Glut1 expression in cells from cluster 0, 7, 11, 12 and all the others. Cells are divided by time point. **(h)** Quantification of Glut1 levels in INFU at 8w pw1, at different distances from the wound site (ratio to 0w is plotted) (left) and whole-mount staining pictures at 0w and 8w pw1 (at ∼ 5 mm from wound site) (right) are shown. n=3-5 mice. **(i)** Time lapse migration assay. Track displacement (μm) of 8w pw1 Glut1^+^ vs Glut1^-^ Lrig1 GL tdTomato^+^ cells. n=3 female mice. P-value: *** 0.0001 to 0.001, ** 0.001 to 0.01, * 0.01 to 0.05. Mean ± SD is plotted if not differently indicated. Scale bars: 50 μm (e, i).

After gathering the cells of the *Transition Cluster* based on time points and clusters, we confirm the intermediate transcriptional state of Lrig1 GL cells at 8w pw1 in-between 0w and healing phases (1w pw1 and 1w pw2), relative to the expression of genes within each GO term (Fig.4c and Extended Data Fig.4a). Specifically, the wound-primed cells are characterised by a cell state that mostly relies on enhanced metabolism and cell motility (Fig.4c and Extended Data Fig.4a), also confirmed through GSEA analysis (Fig.4d and Extended Data Fig.4b). In addition, we find a similar trend for all the GO terms between cells at 8w pw1 and 0w resident in the *Niche of Origin-JZ Cluster* (Fig.4c and Extended Data Fig.4a), indicating that, beside the primed cells in INFU, some Lrig1^+^ SCs in JZ adapted to wound healing. Consistently with cell motility-related GO terms analysis, the infundibular Lrig1 GL cells display an elongated shape at 8w pw1, when compared to typical cuboidal-shape at 0w (Fig.4e,f).

Strikingly, this memory-associated cellular phenotype, once again, is not restricted to the HFs localised at wound proximity but it is also spatially organised in distal HFs with a gradient pattern (Fig.4f), as previously observed (Fig.3k). Similarly, mTORC1 Signalling GO term have been validated as well, through phosphor-S6 (S235-236) staining in distal INFUs (Extended Data Fig.4c).

Concerning the metabolic state, Hypoxia, Glycolysis and Oxidative phosphorylation GO terms are significantly enriched in *Transition cluster* of second wound healing, suggesting an enhanced metabolic state. The glycolytic state is assessed evaluating the levels of Glut1 (Slc2a1), a marker of highly glycolytic cells^27^, that is upregulated specifically in the *Transition Cluster* in actively repairing cells (Fig.4g and Extended Data Fig.4d) and strongly induced at 1w pw2 compared to 1w pw1 (Fig.4g and Extended Data Fig.4e). Flow cytometry quantification validates the Glut1 modulation in our two consecutive injuries model highlighting the increased number of Glut1-expressing Lrig1 GL cells at 8w pw1 when compared to time 0w (Extended Data Fig.4f,g). MitoTracker is used to validate Oxidative Phosphorylation GO term and confirms the scRNA-Seq data (Extended Data Fig.4h). These data support the enhanced metabolism scenario.

From a spatial point of view, high Glut1 expression and oxidative phosphorylation, as markers of primed cells at 8w pw1, display both a gradient pattern with high signal closer to the wound and a weaker signal further away, until ∼ 7 mm from wound edge (Fig.4h and Extended Data Fig.4h). This pattern matches the other observations on krt6, cell shape changes and pS6 (Fig.3k, Fig.4f and Extended Data Fig.4c), corroborating the spatial extent of wound-elicited cell education.

Finally, the enhanced migratory potential of isolated distal memory cells (Glut1^+^TdTomato^+^) relative to non-memory counterpart (Glut1^-^TdTomato^+^) represents a clear evidence of the functional implications that distal priming has on cell fitness (Fig.4i and Extended Data Fig.4i).

Overall, the integration of scRNA-Seq with histological validations allows a deep characterisation of wound-primed cells resident in distal HFs, that relies on high metabolism and enhanced migration rate. Strikingly the histological analysis of the molecular signature of *Priming* (including Krt6a in Fig.3j,k, cell elongation in Fig.4f and Glut1 expression in Fig.4h) shows a spatial distribution of pre-activated Lrig1^+^ GL cells in the INFU that gradually decreases when distance from the wound increase, corroborating the existence of an unexpected large spatial extent of *wound priming*.

### Epithelial priming is lineage specific

Bulge SCs have been shown to exhibit wound adaptation after becoming resident in the repaired niche^15^. Our bulk RNA-Seq analysis on Lgr5 GL cells and the histological data confirm that Lgr5 progeny could be wound-trained even in its own niche, as Lrig1 GL cells do, but with an individual transcriptional program (Fig.1c and Fig.2a). To dissect the adaptation program of Lgr5^+^ bulge SC progeny we performed longitudinal scRNA-Seq on Lgr5 GL cells. Importantly, as evident by histology and scRNA-Seq data (Extended Data Fig.5a-d), no Lgr5 GL cells occupy the INFU when homeostasis is re-established at 8w pw1, demonstrating that only activated Lrig1^+^ SCs give rise to the infundibular primed cells. The *Transition Cluster* contains actively repairing Lgr5 GL cells (1w pw1 and 1w pw2) that are proliferative (Extended Data Fig.5e) and Glut1 and Krt6a positive (Extended Data Fig.5f), as for Lrig1 lineage. Targeted analysis of the GO terms enriched in Lrig1 GL cells reveal that Lgr5 GL cells at 8w pw1 are transcriptionally comparable to cells at 0w (Extended Data Fig.5g), suggesting the absence of a substantial cell priming.

To directly compare the two lineages, we performed pseudotime analysis on Lgr5 GL cells where cluster 5 (the *Niche of Origin-Bulge Cluster*) is used as the starting point of the trajectory D (Extended Data Fig.5h). Focusing on cell plasticity genes in the two D trajectories we noticed that: (i) priming of Lgr5 SCs in their niche of origin is extremely limited compared to Lrig1^+^ SCs (cell subset “A” in Figure S5H), suggesting that the bulge-derived progeny adapt through a trained memory strategy as previously demonstrated^15^; (ii) activated Lgr5 GL cells at 1w pw2 has a smaller enhanced transcriptional activation in comparison with Lrig1 GL cells, consistently with their lower contribution to repair (Fig.1c) (cell subset “B” in Extended Data Fig.5h); (iii) only Lrig1 GL cells has a primed adaptive memory after first injury while establishing an INFU progeny, long after injury resolution and still constitutively express cell plasticity genes (cell subset “C” in Extended Data Fig.5h). Interestingly, although the GO terms enriched for cell plasticity genes are similar in the two linages (Extended Data Fig.5i), the individual genes are mostly particular to each lineage (Extended Data Fig.5j). Thus, we conclude that, in the context of skin full-thickness wound healing, the Lgr5^+^ SCs display a trained adaptation to wound, consistently to what was previously shown in the context of intermediate wound^15^. However, primed cells are peculiar to the Lrig1 SC progeny.

### Long term maintenance of memory through wound-primed progenitors

It has been shown that in epidermal cells wound memory can last up to 80 days^15^. To evaluate how long the wound-distal primed cells are preserved during ageing, we analyse the epidermis of 1.5-year-old Lrig1 GL mice, 10 months after the first injury is induced. The second healing process on the previously injured skin is compared to a first wound repair in aged match mice (Fig.5a). A closure advantage at 1w pw2(40) respect 1w pw1(40), is evident (Fig.5b,c). Importantly even in these aged skin settings, we observe the HF engagement phenotype (Fig.5d) as well as the ex vivo enhanced migratory ability for wound-distal Lrig1 GL cells (Fig.5e), suggesting that distal memory cells are maintained.

**Fig.5:**
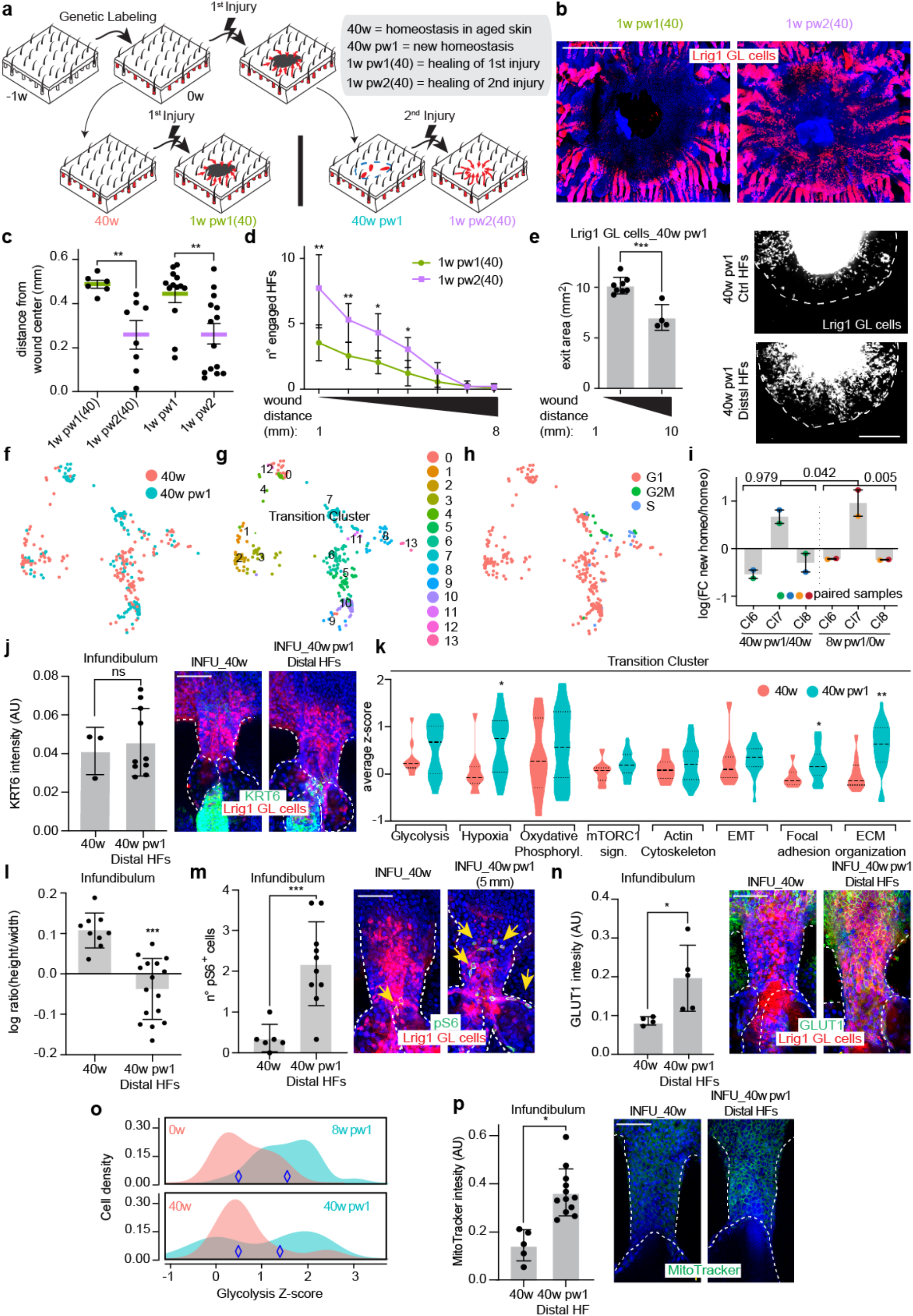
Wound-distal primed cells are preserved in aging. **(a)** Setting to evaluate long term wound memory in Lrig1 GL skin: (−1w) genetic labelling; (0w) wounded and unwounded mice; 40 weeks after labelling, full thickness biopsy on both unwounded (40w) and wounded (40w pw1) mice; 1 week after wound samples are collected, 1w pw1(40) or 1w pw2(40). **(b, c)** Epidermal whole-mount showing Lrig1 GL tdTomato^+^ cells at wound site at 1w pw1(40) and 1w pw2(40) time points (b) and quantification of distance from the wound center (c). Results are compared to young samples at the same time points. n=6-14 wounds. **(d)** Number of the engaged HFs at 1w pw1(40) and 1w pw2(40) from 1 to 8 mm from wound edge. n=6-7 wounds. **(e)** Ex vivo migration assay. Left panel: quantification of the exit area (mm^2^) of Lrig1 GL cells from explants collected from distal memory HFs (within 5 mm) or in Ctrl HFs (more than 10 mm). Right panel: stereomicroscope images of Lrig1 GL tdTomato^+^ cells migration from skin explant. n=4-8 explants. **(f, g, h)** scRNA-Seq of Lrig1 GL cells at the two aged homeostasis: 40w and 40w pw1 time points. UMAP of cells coloured by time points (f), clusters (g) or cell-cycle phases are reported (h). **(i)** Comparison of cell lineages derived from Lrig1 stem cells in two old (40w and 40w pw1, ∼ 5 mm from wound site) and two young (0w and 8w pw1) homeostasis. Ratio from homeostasis (homeo) and new homeostasis cells is plotted. **(j)** Krt6 is quantified (left) in the infundibulum (INFU) from epidermal whole-mount staining (right) at 40w or 40w pw1 (∼ 5 mm from wound site) in Lrig1 GL cells. n=3-10 wounds. **(k)** Violin plot of the average expression for each GO term enriched in young samples comparing 40w and 40w pw1. **(l, m, n)** Cell shape (l), pS6 (m) and Glut1 (n) are quantified (left) in INFU from epidermal whole-mount staining (right) at 40w or 40w pw1 (∼ 5 mm from wound site) in Lrig1 GL cells. Cell shape is reported as ratio between height and width (n=9-16 wounds). Number of pS6^+^ cells is plotted (n=6-10 wounds). Glut1 is stained in n=4-5 wounds. **(o)** Density plot for glycolysis gene signature in Lrig1 GL cells from *Transition Cluster* is plotted comparing the two old (40w and 40w pw1) and two young (0w and 8w pw1) homeostasis. **(p)** MitoTracker levels are quantified (left) in the INFU from epidermal whole-mount staining (right) at 40w or 40w pw1 (∼ 5 mm from wound site). n=5-12 wounds. P-value: *** 0.0001 to 0.001, ** 0.001 to 0.01, * 0.01 to 0.05. Mean ± SD is plotted if not differently indicated. Scale bars 1 mm (b, e). Scale bars: 50 μm (j, m, n, p).

To confirm this, we performed scRNA-Seq of homeostatic skin of old mice (40w) and compare to new homeostasis at 40w pw1 (Fig.5f-h and Extended Data Fig.5k). Strikingly, ∼1 year after injury the infundibular primed cells are well maintained in the *Transition Cluster*, respect to other cell differentiated lineages derived from Lrig1^+^ SCs, such as cells in cluster 6 and cluster 8 (Fig.5i). However, while Lrig1 SC progeny in the INFU does not express krt6, thus suggesting that part of the memory might be lost (Fig.5j and Extended Data Fig.5l), the GO terms enriched in the *Transition Cluster* in young 8w pw1 still follow an overexpression trend ∼1 year after wound when compared to the aged match unwounded (Fig.5k). To validate the GO analysis findings, we performed whole mount histological analysis, as in young skin. Comparing the homeostatic cells at 40w pw1 located millimetres away from the injury with the homeostatic INFU cells at 40w, we still observe cell elongation and increased level of pS6, Glut1 and Mitotraker (Fig.5l-p).

Thus, we demonstrate that the wound memory is largely stable, functional and maintained by wound-distal primed progenitors that are preserved in aged skin for a remarkably long-time frame.

### Long-range priming relies on H2AK119ub-mediated transcriptional de-repression

Chromatin accessibility assays already proved that wound memory involves chromatin changes in trained epidermal cells^15,28^. Even if some transcription factors, functionally determinant in inflammatory memory^12^(Larsen et al., 2021), are likely to be functional also in wound memory, which are the epigenetic regulators is still unknown.

To this aim, we performed an in vivo epigenetic drug screening in Lrig1 GL mice to target five transcriptionally relevant histone modifications, trying to modulate wound-primed cells at 8w pw1. Pre-treatment with PRT4165 does not only lead to a more efficient healing at 1w pw2 (Extended Data Fig.6a,b) but, more importantly, to an increased engagement of HFs located distally from the injury site (Extended Data Fig.6b,c). This drug inhibits the enzymatic activity of Ring1a/Ring1b, component of the Polycomb repressive complexes 1 (PRC1), responsible for the monoubiquitination of lysin 119 on histone H2A (H2AK119ub)^30–32^. To verify the physiological role of PRC1 in *wound priming*, we stained whole-mount epidermis for H2AK119ub at 0w, 1w pw1, 8w pw1 and 1w pw2. In homeostasis, the JZ and the INFU already express lower levels of this histone modification, when compared to other epidermal compartments (Extended Data Fig.6d). When injury occurs and chromatin becomes more accessible^12^, H2AK119ub repressive mark decreases in INFU during both wound healing time points (1w pw1 and 1w pw2) (Fig.6a). When the new homeostasis is re-established (8w pw1), the H2AK119ub mark never restores the original 0w levels (Fig.6a), not even 40 weeks post-wound (aged new homeostasis) (Extended Data Fig.6e). Importantly, the decrease in H2AK119ub is also evident in the INFU of wound-distal HFs (Fig.6a). Since the reduction of H2AK119ub follows the spatial distribution of wound-primed cells (Fig.3 and Fig.4) we hypothesised H2AK119ub to be a key functional component in distal priming through a transcriptional de-repression mechanism. To verify it, after genetical labelling, we pre-treated 0w epidermis with PRT4165 (PRC1i) and we induced injury. PRC1i indeed elicits an enhanced healing rate by Lrig1 GL progeny (Fig.6b-d). Strikingly, wound-distal HFs are also engaged (Extended Data Fig.6f) although in a less extent in comparison 1w pw2 (Fig.1c,d).

**Fig.6:**
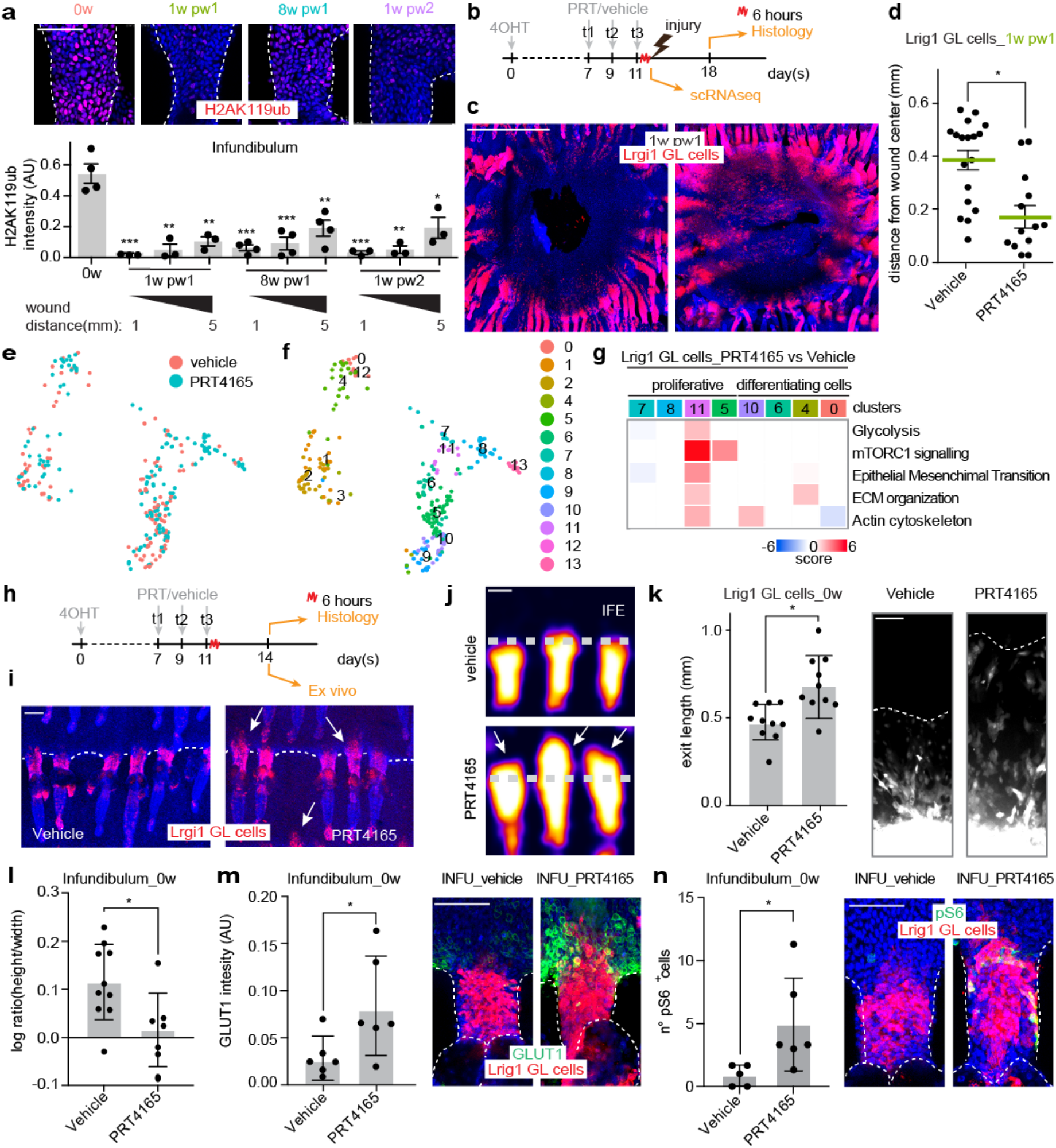
Transcriptional de-repression is a functional component of wound memory. **(a)** Representative pictures (up) and quantifications (down) of H2AK119 monoubiquitination (H2AK119ub) in infundibulum (INFU) at 0w, 8w pw1, 1w pw1 and 1w pw2, at increasing distances from wound site. n=3-4 wounds. **(b)** Setting for PRT4165 treatment in Lrig1 GL mice: (day 0) skin is genetically labelled; (day 7) PRT4165/ vehicle treatment every other day for 3 times; (day7) 6h after the last treatment scRNA-Seq or injury induction; (day 14) at 1w pw1, histology. **(c, d)** Epidermal whole-mount showing Lrig1 GL tdTomato^+^ cells at wound site (scale bar 1 mm) (c) and quantification of distance from the wound center (d) in vehicle vs PRT4165 treated samples at 1w pw1. Mean with SEM is plotted. n=13-18 wounds. **(e, f)** scRNA-Seq data from Lrig1 GL cells treated or not with PRT41645. UMAP of cells coloured by treatment (e) or cluster (f). **(g)** Heatmap of GO terms from Fig.4c, showing gene expression ratio between PRT4165 treated and vehicle cells, per cluster. **(h)** As in **(a)** except for collection day, 3 days after the last treatment. **(i)** Confocal pictures of Lrig1 GL tdTomato^+^ cells vehicle vs PRT4165 treated, 3 days after the last treatment. **(j)** 3D surface plot of Lrig1 GL cells with vehicle/PRT4165 treatment. Dashed line represents IFE-HF boundary. **(k)** Quantification (left) and representative pictures (right) of ex vivo migration assay on vehicle/PRT4165 treated Lrig1 GL cells. n=10 explants. **(l, m, n)** Cell shape (m), Glut1 (n) and pS6 (o) are quantified (left) in INFU from whole-mount staining (right) of epidermis vehicle/PRT4165 treated. Cell shape is reported as ratio between height and width (n = 7-10 wounds) (m); Glut1 is stained in n=6 wounds (n); number of pS6^+^ cells in is plotted (n=5-6 wounds) (o). P-value: *** 0.0001 to 0.001, ** 0.001 to 0.01, * 0.01 to 0.05. Mean ± SD is plotted if not differently indicated. Scale bars: 50 μm (a, i, j, k, m, n).

To assess whether PRC1i promotes features of wound memory even at transcriptional level through transcriptional de-repression of genes associated to *wound priming*, we performed a scRNA-Seq on vehicle or PRC1i treated Lrig1 GL mice (Fig.6e,f and Extended Data Fig.6g,h). Even in the absence of an injury, PRC1i activates several pathways previously identified in the physiological memory of primed cells, especially genes related to GO terms such as glycolysis, EMT and actin cytoskeleton (Fig.6g and Extended Data Fig.6i) in Lrig1 GL cells in the *Niche of Origin-JZ Cluster*. Three days post treatment, this triggers Lrig1 SC progeny to move from the JZ niche into INFU toward IFE (Fig.6h-j), through a cell mechanism that involves cell-autonomous enhanced migration (Fig.6k). Similarly, to wound memory cells, Lrig1 GL cells treated with PRC1i, three days post treatment, now reside in the INFU and are more elongated. In addition, they express more Glut1 and pS6 with respect to vehicle treated (Fig.6m-o), even if, in the absence of a lesion, Krt6 activation signal is not present (Extended Data Fig.6j,k).

Conversely to Lrig1 GL cells, PRC1i treatment to Lgr5 or Gata6 GL skin does not lead to either distal HFs engagement or enhanced wound closure compared to the untreated counterpart (Extended Data Fig.6l-n). However, Lgr5 GL cells 3 days after the last PRC1i treatment only exit their niche of origin towards the upper bulge (Extended Data Fig.6l), in accordance with previous observations^33^.

In conclusion, H2AK119ub in the INFU decreases physiologically after the first injury and the original level is not re-established at 8w pw1 new homeostasis following the distal spatial distribution of priming. Our data indicate that the physiological reduction of H2AK119ub is a functional event, leading to a de-repression of primed genes in distal memory cells derived from Lrig1 SCs.

### Epithelial priming promotes tumour onset in a spatial dependent manner

Wound memory has a beneficial impact on long-term tissue fitness related to skin repair (Fig.5). However, since wound healing and cancer share many hallmarks^34^, we hypothesised that *wound priming* might impact on ageing associated diseases such as tumorigenesis, in line with recent observations in the context of pancreatic cancer^16,35^. In addition, as *wound priming* has a long-range spatial distribution, the detrimental effect might follow this spatial trend.

Importantly, while tissue thickening is a consequence of acute UVB-treatment in both 8w pw1 and 0w homeostatic conditions (Extended Data Fig.7a,b), the engagement of Lrig1 GL cells in IFE is specific to UVB-treated cells at 8w pw1 (Extended Data Fig.7c,d), suggesting that the memory of an antecedent lesion exacerbates the Lrig1 GL cells response to an oncogenic stimulus, potentially triggering cancer. To verify these hypotheses, we induced carcinomas formation in mice through chronic UVB treatment (TS), a physiological oncogenic stress for skin pathology^36^ as in the scheme in Extended Data Fig.7e. This protocol allowed to bypass the papilloma formation step, typical of the chemical induced carcinogenesis in murine skin but absent in human skin tumorigenesis^37^. The comparison of wounded and treated (Wd&TS) with treated-only mice (TS) during chronic UVB-treatment reveals the onset of epidermal dysplasia, typical of actinic keratosis or early Squamous Cell Carcinoma in situ (eSCC)^38^ specifically in Wd&TS mice tail skin (Fig.7a,b and Extended Data Fig.7f,g). eSCC derive mainly from Lrig1 GL primed cells (Extended Data Fig.7h). Strikingly, in clear parallelism with the spatial distribution of primed Lrig1 GL cells (Fig.1d-g and Fig.4e,g), eSCC are distributed following a long-range spatial gradient from wound edge (Fig.7b).

**Fig.7:**
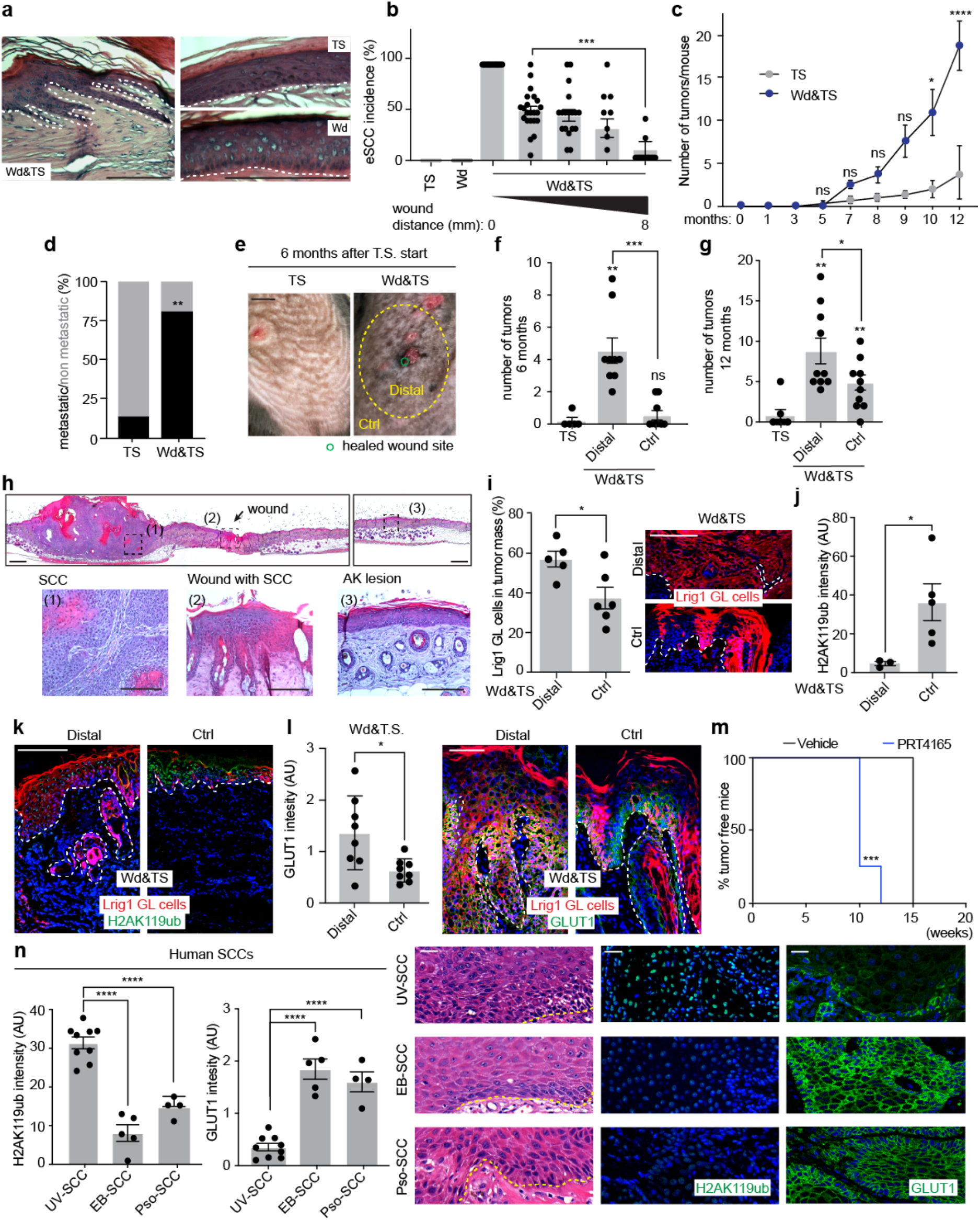
Spatial gradient of wound memory boosted tumorigenesis. **(a, b)** Characterisation of early SCC phenotype in tail skin. Representative pictures (a) and incidence of eSCC is quantified in the indicated samples in a distance dependent manner from wound. Mean with SEM is plotted (b). **(c)** Number of newly formed tumours in the back skin of UV treated mice with a precedent healed wound (Wd&TS) or unwounded (TS, used as controls), shown as a time course. n=3 TS, n=9 Wd&TS Statistics: 2-way ANOVA. **(d)** Presence of lung metastasis was assessed by histology in TS vs Wd&TS. Percentage of metastatic vs not metastatic mice is plotted. n=6 TS, n=8 Wd&TS. Statistics: Chi-square test. **(e)** Representative macroscopic images of the back skin from pre-wounded or control mice. Yellow dashed circles delimit a zone of 1 cm ray around the wound (Distal) that has memory, while outside is control zone (Ctrl) without any memory. **(f, g)** Number of tumours at 6 months (f) or 12 months (g) after tumour stimuli induction (TS) in the indicated conditions. n=7 TS, n=10 Wd&TS **(h)** Tiling (up) and magnification (down) of some selected zones (down), stained with H&E. AK = Actinic keratosis. **(i)** Percentage (left) and representative pictures (right) of Lrig1 GL cells within the tumour mass. n=5 Ctrl, n=6 Distal. **(j, k)** H2AK119ub levels in skin section from Distal or Ctrl zone. Quantification (j) and pictures (k) are shown. n=3 Ctrl, n=5 Distal tumours. **(l)** Glut1 expression in skin section from Distal or Ctrl zone. Quantification (left) and pictures (right) are shown. n=8 tumours/ group. **(m)** Tumour incidence on hairless mice treated with PRT4165. n=7 vehicle, n=4 PRT4165 treated mice. Statistic: Mantel-Cox test. **(n)** H2AK119ub and Glut1 staining on human samples. P-value: **** < 0.0001, *** 0.0001 to 0.001, ** 0.001 to 0.01, * 0.01 to 0.05. Mean with SD is plotted if not differently indicated. Scale bars: 500 μm (a, h(up), i, k); 100 μm (h(down), l, n).

We then verified if the spatial localization of tumours also occurred in back skin, where more severe lesions were expected, as back skin is less resistant to tumorigenesis than tail skin^39^. Consistently with eSCC incidence in tail, Wd&TS back skins show a greater number of cutaneous SCCs compared to TS controls (Fig.7c and Extended Data Fig.7i) with an increased number of metastatic tumours (Fig.7d and extended Data Fig.7j). In addition, the number of formed SCCs (Fig.7e-g), the severity grade of the lesions (Fig.7h and Extended Data Fig.7k,l), as well as the percentage of Lrig1 lineage-derived cells within the tumour mass (Fig.7i and Extended Data Fig.7i) are increased in Wd&T. Remarkably, all these phenotypes have a gradient spatial distribution that overlaps with the location of wound-distal primed cells (Fig.7e-i).

The level of H2AK119ub, whose reduction was functional in distal memory establishment, is lower in SCCs that develop with the contribution of wound-educated Lrig1 GL cells in comparison to tumours with untrained Lrig1 GL cells (Fig.7j,k) and negatively correlates with the overexpression of the priming marker Glut1 (Fig.7l). Moreover, the treatment with PRC1i, which could transcriptionally recapitulate several features of wound priming, accelerate tumour onset in hairless mice (Fig.7m). Overall, long term *wound priming* promotes skin tumorigenesis following its spatial gradient distribution.

Finally, we evaluated the relevance of our findings in the human SCC context, comparing cutaneous SCCs of different causal origins. We focused on Epidermolysis Bullosa derived SCCs (EB-SCCs) and SCCs from psoriatic skin (Pso-SCCs), since these patients have higher risk of tumorigenesis, possibly also through a mechanism that involves respectively wound and inflammatory memory^40,41^. As control we employed SCCs derived from simple UV exposure during life (UV-SCCs). Consistently with our murine observations, Pso-SCCs and EB-SCCs, have a decreased H2AK119ub when compared to UV-SCCs that is associated once again to an enhanced Glut1 expression (Fig.7n), pointing out a memory-tumorigenesis link in humans.

All together, we demonstrate that *wound priming*, widely spatially distributed around the antecedent injury, has long-term detrimental effect on epidermal fitness, enhancing the incidence of squamous cell carcinomas and this is relevant in human context.

## Discussion

### Trained memory versus priming in epithelial cells

Innate immune cells adapt to a stressful event, keep an epigenetic memory of it and respond faster to a second assault through different adaptation programs such as *trained immunity* and *priming*^17,42,43^. The spectrum of epithelial cell responses and their adaptation mechanisms to a stressful event has just started to be understood. *Trained wound-memory* has been reported: HFSCs while repairing an injury, acquire transcriptionally dormant epigenetic memory that can be quickly reactivated upon a subsequent stimulus^15^. Lineage tracing of multiple epithelial lineages combined with scRNA-Seq over two consecutive skin injures allowed us to understand the heterogeneity of adaptive programs other than trained memory. We demonstrated that both Lgr5^+^ and Lrig1^+^ HFSCs acquire wound memory in their respective niche, with different transcriptional programs. However, primed cells are generated only within the Lrig1 progeny during the healing and are maintained in the newly established homeostasis. Strikingly, the transcriptional profile of these cells, intermediate between repairing and homeostatic cells, is a classical feature of *priming* which has not seen before in epithelial context.

### Wide distribution of wound priming in undamaged areas

The spatial extent of wound memory has not previously been assessed due to the absence of an identifiable cell state. Our results demonstrated that the memory has an unprecedent spatial extension that is much larger than expected. In particular, wound-primed progenitors derived from Lrig1 SCs with the described memory cell state, are present up to ∼ 7 mm away from the injury margins. Since the skin is the largest organ in mammals this spatially large adaptation involves tissue at sub-organ scale. However, in other smaller epithelia it is likely that this memory might impact the whole organ.

### Transcriptional de-repression and cell state of wound-primed progenitors

Epigenetic studies of inflammatory memory in epidermal SCs demonstrated increase of chromatin accessibility together with active histone marks deposition^12^. This chromatin scenario is compatible with a reduction of transcriptional repressive histone marks. In particular, we observed a clear correlation between the spatial distribution of decreased levels of H2AK119ub in the INFU after wound and the wide location of the *wound priming* for up to ∼ 7 mm from the injury. Since PRC1 inhibition leads to acquisition of some enhanced repair skills and to the transcriptional de-repression of several gene sets typical of *wound priming* in Lrig1 but not in the Lgr5 progeny, we concluded that the reduction of H2AK119ub functionally contributes to priming only in a specific cell identity context. This, together with the observation that the upper HF is characterised by a low level of H2AK119ub even in homeostatic conditions, conferred to the Lrig1^+^ SCs a new repair-specific role. Thus, it is easy to speculate that specific cells in other epithelia might also have this unique innate potential.

mTOR- and HIF-1α-mediated Glycolysis has been identified as the metabolic basis for *trained immunity*^44^. Interestingly, the cell state of the identified wound-primed cells in the infundibulum is characterised by similar features and enhanced migratory abilities. Since we recapitulated this cell state through PRC1 inhibition, we established a direct causal link between the identified epigenetic and metabolic state. Because all the above-mentioned similarities between the adaptation programs of epithelial and innate immune cells, it is likely that the reduction of H2AK119ub that we identify in epithelial cell priming might be functional in immune cell context.

### Long-term priming impacts skin cancer susceptibility

Beside the positive effect of wound memory in tissue repair context, we demonstrate that skin tumorigenesis is favoured by the presence of a previous completely resolved injury and that Lrig1 GL progeny has a major contribution within the tumour mass. Strikingly, the correlation between the spatial distribution of primed cells and the tumour onset corroborates the link between wound memory and cancer. Mechanistically, our data supported the hypothesis that the reduction of H2AK119ub, a functional feature of *wound priming*, directly promotes tumorigenesis. On this line, wound memory associated tumours have reduced H2AK119ub levels in comparison to control tumours, not only in mice but also in humans. Our results together with recent findings in diversified cellular contexts^45,46^, supported the general conclusion that loss of key histone repressive mark promotes tumorigenesis. This raises a general concern with respect to the therapeutic intervention on repressive chromatin factors that could be beneficial in regenerative medicine but detrimental in oncology.

## Acknowledgments

We thank Paolo Porporato, Carlos Sebastian, Klaas Mulder and Samuel Woodhouse for their discussions and suggestions. We thank all the facilities at the Molecular Biotechnology Center for their technical support and all the MBC members for making the Institute a friendly and scientifically stimulating environment. The G.D. laboratory is supported by AIRC, Associazione Italiana per la Ricerca sul Cancro (MFAG 2018 - Id.21640) and by CZI, Chan Zuckerberg Initiative, an advised fund of Silicon Valley Community Foundation (DAF2020-217532). MW is a recipient of JSPS overseas research fellowship. The S.O. laboratory is supported by AIRC (IG 2017 - Id. 20240)

## Author Contributions

GD designed and supervised the study. CLL and MW performed all the experiments with CD and DD assistance. Sequencing samples were prepared by VP, sequenced by FA and analysed by GP, AL and SK. TN and KN provided human samples. DB performed flow cytometry analysis. GD, CLL, MW, VP, TH, KN and SO interpreted the data. GD and CLL wrote the manuscript with input from all authors.

## Methods

### Mouse strains

Maintenance, care and experimental procedures have been approved by the Italian Ministry of Health, in accordance with Italian legislation and the institutional review board of the Hokkaido University Graduate School of Medicine. Rosa26-fl/STOP/fl-tdTomato, Lgr5-EGFP-ires-CreERT2, Gata6-EGFP-ires-CreERT2 and Lrig1-EGFP-ires-CreERT2 have been previously described^11,18,19,48^. For tumorigenesis, SKH-1 hairless mice have been purchased from Charles-River Laboratories. Hairless mice were maintained in specific pathogen free conditions; the other mice were kept in conventional conditions with 12hr light dark cycles from 8 am to 8 pm. The study is compliant with all relevant ethical regulations regarding animal research.

### Lineage tracing of hair follicle lineages

For lineage tracing, Lgr5-EGFP-ires-CreERT2, Gata6-EGFP-ires-CreERT2 or Lrig1-EGFP-ires-CreERT2 were crossed with Rosa26-fl/STOP/fl-tdTomato strain. Genetical labelling was induced in epidermis of 6-8 weeks old mice with a single topical administration of 75 μg of (Z)-4-Hydroxytamoxifen (Sigma-Aldrich) (15 mg/mL diluted in acetone).

### Full thickness wounding

1 week post tamoxifen-treatment (0w), mice were anesthetised with intramuscular injection of Ketamine/Xylazine (Rompun®) 30 μl/head or isoflurane inhalation. Full thickness wounds were made with a circular biopsy punch in tail (2 mm) or dorsal (5 mm) skin. For dorsal skin wounds mice were shaved with an electric clipper 24 h before the procedure.

### Epidermal and dermal whole-mount and immunostaining

Tail epidermal or dermal whole-mounts were prepared as previously described^18^. Briefly, the skin was incubated in PBS/EDTA (5mM) at 37 °C for 4 h to separate the epidermis from the dermis using forceps. The collected epidermis and dermis were pre-fixed 1h in 4% paraformaldehyde (PFA, Sigma-Aldrich), rinsed three times with PBS for 5 min and conserved in PBS with 0.2% azide at 4°C, till usage. For immunofluorescence staining, epidermis/dermis samples were permeabilised and blocked in PB buffer (0.5% skim milk, 0,25% fish gelatin and 1% of Triton X-100 in PBS) for 2 h at room temperature (RT) (for epidermis) and overnight at 4 °C (dermis). Subsequently, primary antibodies were diluted in PB buffer at the indicated concentration (see below) and incubated overnight at 4 °C. The following primary antibodies and dilutions were used: anti-Keratin 6A (rabbit, 1:200, Biolegend_905701), anti-Ubiquityl-Histone H2A (Lys119) (clone D27C4) (rabbit, 1:1000, CST_8240), anti-phospho-S6 Ribosomal Protein (Ser235/236) (clone D57.2.2E) (rabbit, 1:200, CST_4858), anti-Ki67 (rabbit, 1:100, Abcam_16667), anti-Glucose Transporter GLUT1 (SLC2A1) (EPR3915 clone) (rabbit, 1:200, Abcam_ab115730), CD45 (rat, 1:100, eBioscience_30F11), Alexa-647 conjugated Phalloidin A (Thermo-Fisher). After 30 min of PBS wash, samples were incubated with secondary antibodies together with 4’,6-diamidine-20-phenylindole dihydrochloride (DAPI) (Thermo-Fisher, diluted 1ug/mL) 2 h at RT (for epidermis) or overnight at 4 °C (for dermis). Alexa Fluor-568-, 488-, or 647-conjugated secondary antibodies (Life Technologies) were used at 1:1000 dilution. Tissues were finally mounted using Mowiol 4-88 Reagent (Sigma-Aldrich) mounting solution.

### Mitochondrial activity

For quantification of mitochondrial activity, MitoTracker staining (MitoTracker® Deep Red FM_M22426, Thermo-Fisher) was performed in whole-mount epidermis according to the datasheet instructions. Briefly, freshly isolated epidermis was incubated in 50 nM MitoTracker solution in DMEM low calcium (Thermo-Fisher) without serum at 37 °C 10 min. The epidermis was then washed with DMEM (3 washes of 20 min) and fixed 40 min 4% PFA before mounting.

### Histology and immunostaining on sections

For frozen section, back skin or tail tumor samples were pre-fixed 20 min at RT in 4% PFA immediately after dissection, to preserve tdTomato signal. After fixation tissues were washed 30 min in PBS, embedded in OCT Compound (Bio Optica) and kept at -80 °C. Frozen tissue blocks were sectioned (5-7 μm) with a CM3050S Leica cryostat (Leica Mycrosystems). Tissues were washed three times in PBS for 5 min and incubated in PB buffer for 1 h at RT. Primary antibodies were incubated overnight at 4 °C. Sections were rinsed three times in PBS and incubated with secondary antibodies and DAPI (Thermo-Fisher, diluted 1ug/mL) in PB buffer for 1 h at RT. Sections were again washed three times with PBS. The following primary antibodies were used: anti-Ubiquityl-Histone H2A (Lys119) (clone D27C4) (rabbit, 1:1000, CST_8240) and anti-Glucose Transporter GLUT1 (SLC2A1) (clone EPR3915) (rabbit, 1:200, abcam_ab115730). For paraffin embedding, freshly collected samples were closed in blocks and fixed 24 h in 10% NBF (Neutral Buffered Formalin, Sigma-Aldrich) and then transferred to 70% EtOH solution until the embedding. Samples were sectioned (5-10 μm) with a micrometer (Leica Mycrosystems). After deparaffinization and hydration hematoxylin and eosin (H&E) staining or immunofluorescent staining were performed. For immunofluorescent staining, antigen unmasking was performed for 20 min at 98 °C in citrate buffer (pH 6). The sections were then blocked with PB buffer for 1 h at RT. Primary and secondary antibody incubations wer performed overnight at 4°C, and 1h RT respectively, followed by DAPI incubation. The following primary antibodies were used: anti-Ubiquityl-Histone H2A (Lys119) (clone D27C4) (rabbit, 1:1000, CST_8240) and anti-SLC2A1 (GLUT1) (rabbit, 1:100, Sigma-Aldrich_ HPA031345). Slides were mounted with ProLong Glass Antifade Mountant (Invitrogen).

### Immunostaining on ex vivo culture

Skin explants and exiting cells were fixed in 4% paraformaldehyde at RT for 10 min. Permeabilisation and blocking were performed using the staining buffer (1%BSA, 5% Fetal Bovine Serum and 0.3% Triton X-100 in PBS) 1 h at RT. Primary antibodies were incubated at the indicated concentration overnight at 4 °C and secondary antibodies used at 1:1000 concentration 1 h at RT. DAPI nuclear staining was incubated in the dark for 5 min at RT. The following primary antibodies and dilutions were used: anti-Keratin 6A (rabbit, 1:500, Biolegend_905701) 1:500, anti-cytokeratin 14 (clone LL002) (mouse IgG3, 1:500, Invitrogen), GM130 (clone 35) (mouse, 1:400, BD pharmingen).

### Microscopy

Images were acquired with: Confocal: Leica TCS SP5 Tandem Scanner; Leica TCS SP8 Tandem Scanner; Stereoscope: Leica MZ16FA; Optical microscope: Olympus Bx41. Whole-mount and section stainings were examined using a Leica TCS SP5/SP8 confocal microscopes equipped with 20x/40x or 60x immersion objectives (Zeiss). Z-stacks were acquired at 400 Hz with an optimal stack distance and 1024×1024 dpi resolution. Z-stack projections were generated using the LAS AF software package (Leica Microsystems) as max intensity projections. For total wound bed images, the TCS SP5 resonance scanner was used. Four laser lines (405, 488, 561 and 633 nm) were used for near simultaneous excitation of DAPI, Alexa448, tdTomato and Alexa647 fluorophores. H&E staining was acquired using Olympus Bx41/ Leica DM6 microscope equipped with a 4x or 10x objective.

### Immunofluorescence intensity measurements

Digital images were processed and analysed using Fiji (https://imagej.nih.gov/ij/). Fluorophore intensity was measured as Integrated Density (IntD) in the selected ROI. The staining intensity (AU) was calculated as the ratio between the IntD of the antibody signal and the IntD of the DAPI signal in the same ROI. 3D surface plot of hair follicles was obtained using the Fiji function 3D surface plot. 5 hair follicle triples were overlapped for the analysis. Images are shown with the pseudo-colour Fire. For phalloidin staining segmentation we used the MorphoLibJ plugin with the Morphological Segmentation function. The results were displayed as Catchment basins option.

### Lrig1 GL cells isolation and culture

Tail skin was dissected and incubated in trypsin EDTA (Thermo-Fisher, 0.25% in PBS) with the dermis side down overnight at 4 °C. The day after, the epidermis was peeled off from the dermis and chopped with 2 scalpels for 1 min. Isolated cells were cultured in keratinocytes media (low-calcium DMEM (Thermo-Fisher) supplemented with 10% FBS (Sigma-Aldrich), 100 U/ml penicillin-streptomycin (Thermo-Fisher), HCE cocktail consisting of hydrocortisone 0.5 μg/ml (Sigma-Aldrich), insulin 5 μg/ml (Thermo-Fisher), cholera enterotoxin 10^−10^ M (Sigma-Aldrich) and EGF 10 ng/ml (PeproTech) and cultured at 34 °C in a humidified atmosphere, with 8% CO_2_.

### In vitro adhesion and survival assay

Isolated epidermal cells were counted (tdTomato^+^) with FACS Verse, plated on 6 well plate (Corning) and treated 2 h with 4 μg/mL of mitomycin C, to stop proliferation. Cells were counted again 24h and at 72h after adhesion plating. Adhesion was calculated as the ratio of plated cells and counted cells at 24h, while survival as the ratio between cell number at 24h and 72h. Percentage was plotted.

### In vitro time lapse migration assay

For time lapse migration assay, isolated cells were plated in μ-Slide (Ibidi) with keratinocytes media. 24 h after plating dead cells were removed by PBS washes and attached cells were used for microscope acquisition. Plates were maintained in the incubator chamber of a confocal TCS SP5 microscope (Leica Microsystems) under controlled conditions (34 °C, 8% CO2). Images were collected in two different positions for each well and acquired every 15 minutes for 16 h. Cell displacement was tracked automatically using the TrackMate plugin on Fiji with LogDetector settings. Mean track displacement was calculated for each sample and plotted. For the normalised start position graph, we subtracted each point in the track with the coordinates of the starting point.

### Ex vivo migration assay

The ex vivo migration assay was performed as previously described^13^. 2 mm punch biopsies were taken from freshly isolated tail skin. The skin was adhered to the bottom of a 24 well plate (Corning) and cultured in complete keratinocyte media under controlled conditions (34°C, 8% CO_2_) for 4, 7, and 9 days, as indicated.

### Fluorescence-activated cell sorting (FACS)

To obtain single cell suspensions for FACS, roughly 5×5 mm of tail skin from 0w, 1w pw1, 2w pw1, 4w pw1, 8w pw1 and 1w pw2 were dissected and incubated in trypsin EDTA (Thermo-Fisher, 0.25% in PBS) with the dermis side down overnight at 4 °C. The day after, the epidermis was peeled off from the dermis with a scalpel and gently minced with 2 scalpels for 1 min. The resulting material was well resuspended with keratinocyte media and then filtered through a 70 μm cell strainer (VWR, Corning). The cell suspension was then centrifuged for 7 min at 4 °C 1500 rpm and washed twice with PBS. For single cell RNA-Seq and mini-bulk RNA-Seq, cells were resuspended in 500 μL of 1% BSA in PBS and 2 μL of SUPERase RNase Inhibitor (Thermo-Fisher, 20 U/μL) was added to each sample. td-Tomato positive cells were sorted with a 100 μm nozzle into 96 well PCR plate (Bio-Rad) containing lysis buffer (0.5% Triton X-100 in H2O). For tdTomato^+^ GLUT1^+^ (SLC2A1) cells sorting, cells were blocked with 3% FBS in PBS for 20 min and then incubated 30 min with 0.3 ug of anti-GLUT1 (clone EPR3915) (Alexa Fluor 647 conjugated, Abcam_ab195020) on ice. Antibody solution was removed, cells were washed three times with cold PBS and sorted in a new collection tube containing keratinocytes media with a 100 μm nozzle. To select viable cells TO-PRO-3 Iodide (Thermo-Fisher) was added at 1 μM concentration to the cell suspension before sorting. BD FACSAria II equipped with Diva software was used (BD Biosciences).

### GLUT1 quantification by flow cytometry

Flow cytometry quantification of GLUT1 (SLC2A1) expression in Lrig1 GL tdTomato+ cells was performed on tail skin samples at different time points. Samples were processed as for sorting, with minor modifications. Briefly, after chopping, the cell suspensions were moved to a V bottom 96 well plate (Corning) for blocking and staining. 1μg/mL DAPI was used to assess cell viability. Single cell suspensions were first gated against DAPI to exclude dead cells, and then with forward and side scatters to isolate singlets. Flow cytometry plots were generated using FlowJo™ v10.8. Manual compensation was performed.

### Experiments with epigenetic drugs

For drug screening 5 drugs targeting epigenetic factors have been used to treat 8w pw1 mice. Briefly drugs were dissolved in acetone at the proper concentration and 50 μL of each drug was applied every other day for three times on tail skin. 6 h after the last treatment Lrig1 GL mice were wounded as described above. 1 week after the second injury (1w pw2) samples were collected and the migration of Lrig1 GL cells migration towards the wound center analysed. Drug list: A-196, SUV420H1 and SUV420H2 inhibitor (Sigma-Aldrich, 3 mg/mL), UNC0638, EHMT1/2 inhibitor (Sigma-Aldrich, 3 mg/mL), SB747651A, MSK1 inhibitor (Axon Medchem, 1.5 mg/mL), EX-527, Sirtuins inhibitor (Sigma-Aldrich, 3 mg/mL), PRT4165, Ring1a/1b inhibitor (PRC1) (Sigma-Aldrich, 3 mg/mL). To test the ability of PRT4165 to mimic memory the same approach reported above was used to treat 0w mice.

### UV irradiation and tumorigenesis

The UVB irradiation protocol was performed as previously described^49,50^. The UVB light source used was UVM-28EL (UVP Ultraviolet Product, Thermo-Fisher). The irradiated energy was regularly measured with PCE-UV36 UVC radiation meter (PCE inst.). For acute UV irradiation, mice were treated with 200 mJ/cm2 for three times consecutively. The samples were collected 2 days after the last irradiation and analysed for epidermal thickness or hair follicle engagement. For in vivo tumorigenesis the DMBA-UVB two-stage-induced carcinogenesis protocol was used. Two days before the DMBA application, mice back skin were shaved with an electric shaver. Mouse dorsal or tail skin was treated with 50 μL of 120 μg/mL of DMBA (Sigma-Aldrich) dissolved in acetone after the complete healing of wound or without wound as control. UVB irradiation (180mJ/cm2) was started 10 days after the DMBA application and continued for 3 times a week till the end of the experiments. All the mice were shaved once every 10 days. At the end points samples were collected, embedded in OCT or paraffin and analysed for histology. The tumours were classified as Squamous cell carcinoma (SCC) in situ or SCC according to the tumour architecture and the cytoplasmic morphology as previously described^49,50^. For tumorigenesis in SKH-1 hairless strain, mice were treated with 50μL of PRT4165 (3 mg/ml) (Sigma-Aldrich) in acetone or with acetone alone every other day for 3 times, prior to UVB irradiation was started. All the mice were irradiated with 250 mJ/cm2, 3 times a week until the end point^51^.

### Human squamous cell carcinomas (SCC)

Cutaneous SCC samples with AK (UV-SCC) regions were obtained from 9 patients. In addition, cutaneous SCC samples were collected from 3 Recessive Dystrophic Epidermolysis Bullosa (RDEB) (EB-SCC) (5 SCCs) and 4 psoriasis patients (Pso-SCC). The institutional review board of the Hokkaido University Graduate School of Medicine approved the human study described (ID: 13-043, 14-063, and 15-029). The study was carried out according to the Declaration of Helsinki Principles. The participants provided written informed consent. The RDEB patients harbored compound heterozygous mutations in COL7A1 (NM_000094.4) (patient 1: c.5443G>A (p.Gly1815Arg) and c.5819del (p.Pro1940Argfs*65), patient 2: c.5932C>T (p.Arg1978*) and c.8029G>A (p.Gly2677Ser), patient 3: c.7723G>A (p.Gly2575Arg) and c.8569G>T (p.Glu2857*)) and the expression of type VII collagen was reduced in their skin specimens^52,53^.

### Tumour analysis

For quantification of effraction of basement membrane zone as indication of tumour invasiveness, the length of deepest site of tumour from the normal adjacent basement membrane zone in H&E staining was evaluated by Fiji. As to grading SCC about differentiation, the ratio of well structure maintained epidermal layers and aberrant epidermal layers in H&E staining are calculated using Fiji.

### Mini-bulk RNA-Seq and analysis

Mini-bulk RNA-Seq have been performed on 150 cells upon polyadenylated mRNA isolation. Briefly, the Smart-Seq2 biotinylated Oligo(dT) was bound to Dynabeads MyOne Streptavidin T1 beads (Thermo-Fisher) and used to isolate mRNA from the cell lysate. Reverse transcription, amplification and library preparation were performed as for the single cell library preparation doubling the volumes. After quality controls with FastQC v0.11.2 (https://www.bioinformatics.babraham.ac.uk/projects/fastqc), raw reads were processed with Trim Galore! v0.5.0 to perform quality and adapter trimming (parameters: --stringency 3 –q 20) (https://www.bioinformatics.babraham.ac.uk/projects/trim_galore). Trimmed reads were next aligned to the mouse reference genome (UCSC mm10/GRCm38) using HiSat2 v2.2.01 with options -N 1 -L 20 -i S,1,0.5 -D 25 -R 5 --pen-noncansplice 20 --mp 1,0 --sp 3,0 and providing a list of known splice sites^54^. Gene expression levels were quantified with featureCounts v1.6.12 (options: -t exon -g gene_name) using the GENCODE (https://www.gencodegenes.org) release M20 annotation^55^. Gene expression counts of each cell population were next analysed using the edgeR3 R/Bioconductor package^56^. Lowly expressed/non-detected genes were filtered out (i.e.1 CPM in less than 3 samples) and normalisation factors were calculated using the trimmed-mean of M-values (TMM) method. In order to account for unwanted sources of noise in the dataset, the normalised counts were processed using the SVA4 R/Bioconductor package^57^. The estimated surrogate variables were then included as covariates in the generalised linear model (GLM) used for differential expression testing (formula: ∼time+sv1+…+svn). Following dispersion estimation, an ANOVA-like test was performed in order to identify the genes that were differentially expressed in any sample group across the time course, using 0w as the reference group in the model. Genes with |logFC|>1 in at least one time point and FDR<0.05 were considered as significantly differentially expressed. The same analyses were performed for each cell population. Memory gene list are defined as the gene with 1w pw1 |logFC| < 1w pw2 |logFC|. GO analysis was performed using EnrichR tool (v 3.0) (https://maayanlab.cloud/Enrichr)^58^.

### scRNA-Sequencing

Epidermal cells were isolated and sorted as previously described. Female mice only were used for this experiment. scRNA-Seq was performed with a modified version of the Smart-Seq2 protocol^59^ as in^60^. Briefly, individual cells are sorted into 96 well plate containing 2 µl of lysis buffer (0.5% Triton X-100 in H_2_O), in presence of RNase inhibitor (Thermo-Fisher), dNTPs and Oligo(dT). Reverse transcription of the polyadenylated RNA has been performed with Maxima Rt Minus Reverse Transcriptase (Thermo-Fisher) and Template Switching Oligos. The resulting cDNA was amplified with 25 cycles of PCR and libraries will be prepared for sequencing with standard NexteraXT Illumina protocol (Illumina).

### scRNA-Sequencing analysis

Reads were mapped to the *Mus musculus* transcriptome (ENSEMBL version 101) using Salmon v 0.13.1. Data processing: cells with less than 1000 genes expressed or with over 50 % of mitochondrial reads were filtered out, genes expressed in less than 5 cells were filtered out. Replicates were normalised using Seurat^61^ and corrected for batch effects using Cluster Similarity Spectrum, Harmony or FastMNN. Optimal clustering was selected based on cumulative gene expression patterns of marker genes. RNA-Seq data were analysed using Seurat packages (v 4.0.1) on R (v 4.0.3). Only protein-coding genes were considered for downstream analysis and read counts selected were batch corrected with the CSS method. All Seurat analyses were performed using default parameters. The specific changes are reported when present. Dimensionality reduction (PCA + UMAP) were calculated with Seurat functions RunPCA (npcs = 100) and RunUMAP (reduction = “css”, dims = 1:8), using the top 10000 variable genes. Cell clustering was performed with function FindNeighbors (reduction = “css”, dims = 1:8) and FindClusters (resolution = 2.1).

Cluster marker genes were identified by FindAllMarkers (only.pos = T, return.thresh = 0.05) function looking for upregulated genes in each cluster against the remaining cells and the top 50 by the “avg_logFC” rank was selected.

### Differential Expression Analysis and feature Extraction

The analysis of differentially expressed genes within the same cluster was performed using Seurat’s FindMarkers() function. To plot the expression of genes or gene groups the FeaturePlot() function.

### Seurat Integration

Single cell data from Lgr5 GL cells, Lrig1 GL cells aged homeostasis or Lrig1 GL cells treated with vehicle or PRT4165 were projected onto the reference UMAP structure derived from Lrig1 GL young data, in order to localise the position of each cell in the hair follicle or interfollicular epidermis. The overlap was performed using SeuratIntegration Tools^62^ with FindTransferAnchors and MapQuery functions.

### Pseudotime analysis

Pseudotime analysis was carried out using Slingshot package (v. 1.8.0)^63^ on the Seurat object, using the top 10000 variable genes. For the pseudotime A, B, C, D, E cluster 11 corresponding to junctional zone stem cells on the basis of the molecular markers, was selected as the starting point. Pseudotime F was instead found selecting cluster 5 (bulge) as starting position. To derive the differentially expressed genes (DEG) the TestPseudotime function was used with a threshold of FDR < 0.05. The genes were z-scored (and smoothed) and uploaded on Morpheus (https://software.broadinstitute.org/morpheus) to obtain the heatmaps. For pseudotime D specifically a deep analysis was performed. 0w and 1w pw1 cells were separated from 8w pw1 and 1w pw2 cells and ordered on pseudotime. Morpheus clustering was then performed (hierarchical clustering, row, complete) on 0w - 1w pw1 cells to compare the expression and the trend of the same genes in the two conditions.

### GO Enrichment

Gene Ontology (GO) enrichment analysis was performed using EnrichR tool (v 3.0)^58^ using GO Biological Process. Results were used to generate “DotHeatmap’’ to compare enrichment between different conditions and pseudotimes.

### GSEA Analysis

GSEA Analysis was performed using the GSEA function by ClusterProfiler package (v 3.18.1)^64^ and using as Database reference: MSigDB (all categories of DB), GO (Biological Process, Molecular Function and Cell Component) and KEGG. DEG genes obtained from 0w vs 8w pw1 and 1w pw1 vs 1w pw2 comparisons were used as input.

### Quantification and statistical analysis

Statistical analysis was performed by using the GraphPad Prism 7 software (GraphPad Inc., San Diego, CA, USA). Data are presented as mean ± standard error of the mean or mean ± standard deviation. Statistical significance for all wound healing studies was determined with the two-tailed unpaired Student’s t-test with a 95% confidence interval under the untested assumption of normality. Statistical significance tumor incidence was calculated using either a Mann-Whitney test or random permutation. No statistical method was applied to predetermine sample size and mice were assigned at random to groups. Images are representative of at least three independent experiments and mice from both sexes were used, if not differently indicated.

## Materials availability

This study did not generate new unique reagents. Material that can be shared will be released via a Material Transfer Agreement. Further information and requests for resources and reagents should be directed to and will be fulfilled by the lead contact: Giacomo Donati at.

## Data and code availability

Data reported in this paper will be shared by the lead contact upon request. This paper does not report original code. Any additional information required to reanalyse the data reported in this paper is available from the lead contact upon request. scRNA-Seq data that support the findings of this study will be deposited in the Gene Expression Omnibus (GEO).

