## Supplementary Figures for "Lifelong tissue memory relies on spatially organised dedicated progenitors located distally from the injury"

Extended Data Fig.1: Characterisation of two consecutive injuries model.

Extended Data Fig.2: Transcriptome of Lrig1 GL cells suggests a new primed subpopulation arising after wound resolution

Extended Data Fig.3: Single cell analysis assigned known epidermal niches to cell clusters

Extended Data Fig.4: Characterisation of primed-memory cell state.

Extended Data Fig.5: Comparison between single cell profiling of Lgr5 and Lrig1 lineages

Extended Data Fig.6: Epigenetic characterisation and manipulation of wound-primed memory.

Extended Data Fig.7: Healed skin is more sensitive to tumour onset

**Extended Data Fig.1**

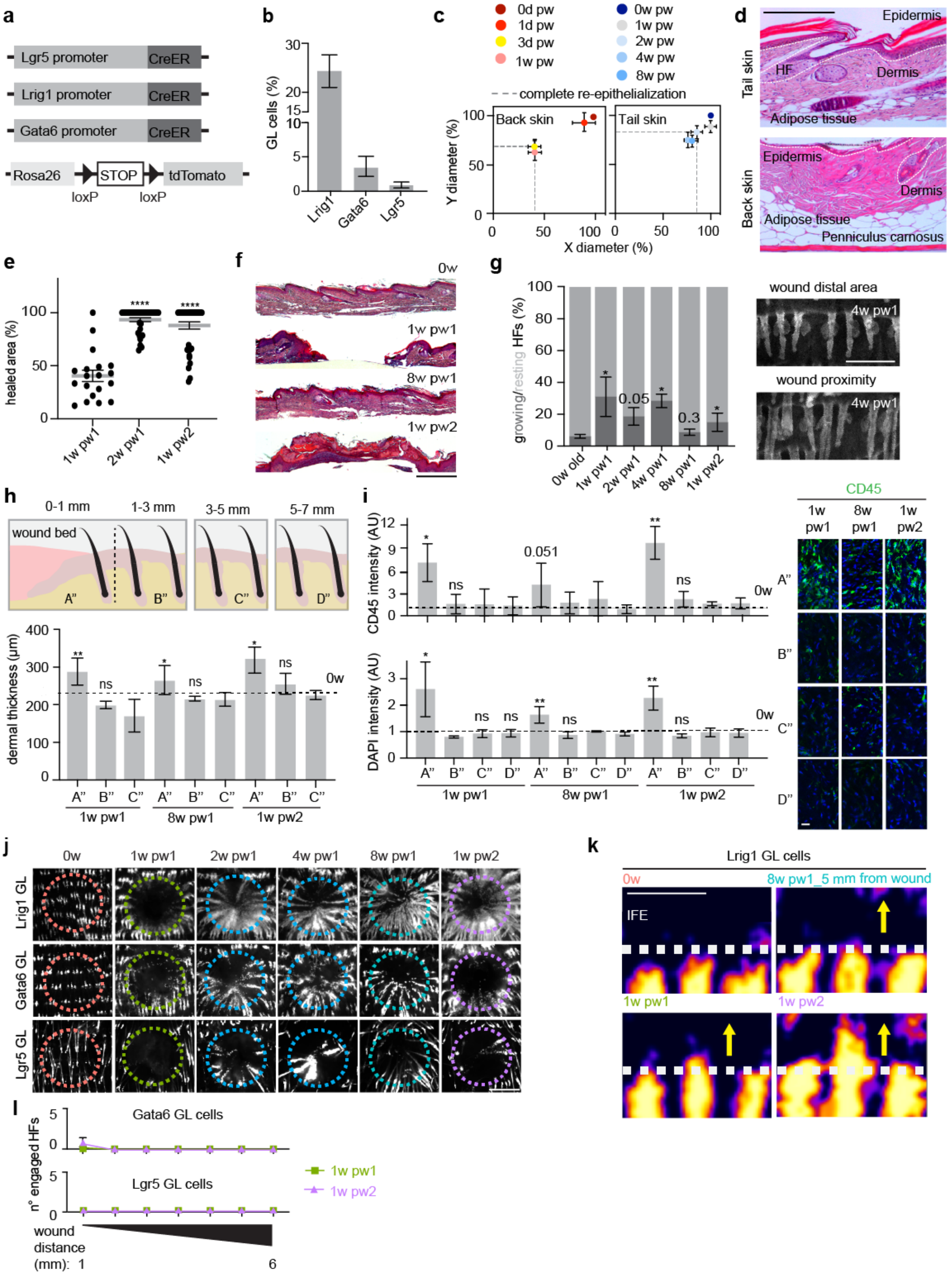

**Extended Data Fig.1: Characterisation of two consecutive injuries model. (a)** Scheme of murine genetic models: Cre-ER is expressed under Lrig1, Lgr5 or Gata6 promoters. Each strain is crossed with Rosa126-STOP-tdTomato mice to achieve genetic labelling upon Cre-ER activation by tamoxifen. **(b)** Flow cytometry quantification of GL cells in the whole epidermis at 0w. n=3-4 mice. **(c, d)** Comparison of wound margins contraction during wound healing in back vs tail (c). Haematoxylin-Eosin (H&E) staining of tail and back murine skin show the absence of panniculus carnosus in tail skin. Dashed line indicates epidermis (d). Tail skin wound model was selected to reduce the issue of tissue contraction. n=2-6 mice. **(e)** Percentage of re-epithelialization area at 1w pw1, 2w pw1 and 1w pw2. n=19-37 wounds. Mean with SEM is plotted. **(f)** H&E staining of skin at indicated time points. **(g)** Evaluation of hair follicle cycle phase. Percentage of resting or growing HFs at different time points (left panel) and whole-mount of tail epidermis at 4w pw1 (right panel) show comparable a HF cycle state between the original (0w) and the newly established (8w pw1) homeostasis. **(h, i)** High cell density, typical of fibrosis, and immune cells infiltration is resolved at 8w pw1. **(h)** Upper panel: Scheme of the sub-regions identified in injured skin; lower panel: quantification of dermal thickness (µm), in A", B" and C" regions. n=3-4 wounds. **(i)** Left panels: CD45 index (CD45 positive cells per $mm^2$) and DAPI index (number of nuclei per $mm^2$) relative to 0w are measured from dermal whole-mounts immunostaining. Mean with SEM is plotted. n=3 mice; right panels: pictures of CD45 staining in the indicated sub-regions. **(j)** Epidermal whole-mounts (red channel) of GL tdTomato$^+$ cells showing their contribution on the wound bed. Dashed circles highlight the original wound perimeter. **(k)** 3D surface plot (red channel only) show GL cells exit from HFs at 1w pw2. Dashed line represents IFE-HF boundary. (l) Number of engaged HFs in Gata6 and Lgr5 GL cells 1w pw1 and 1w pw2 up to 6 mm from wound edge. n=8-10 wounds. P-value: **** < 0.0001, ** 0.001 to 0.01, * 0.01 to 0.05, ns ≥ 0.05. Mean with SD is plotted if not differently indicated. Scale bars: 300 µm (d, f, g); 50 µm (i); 1 mm (j, k).

### Extended Data Fig.2

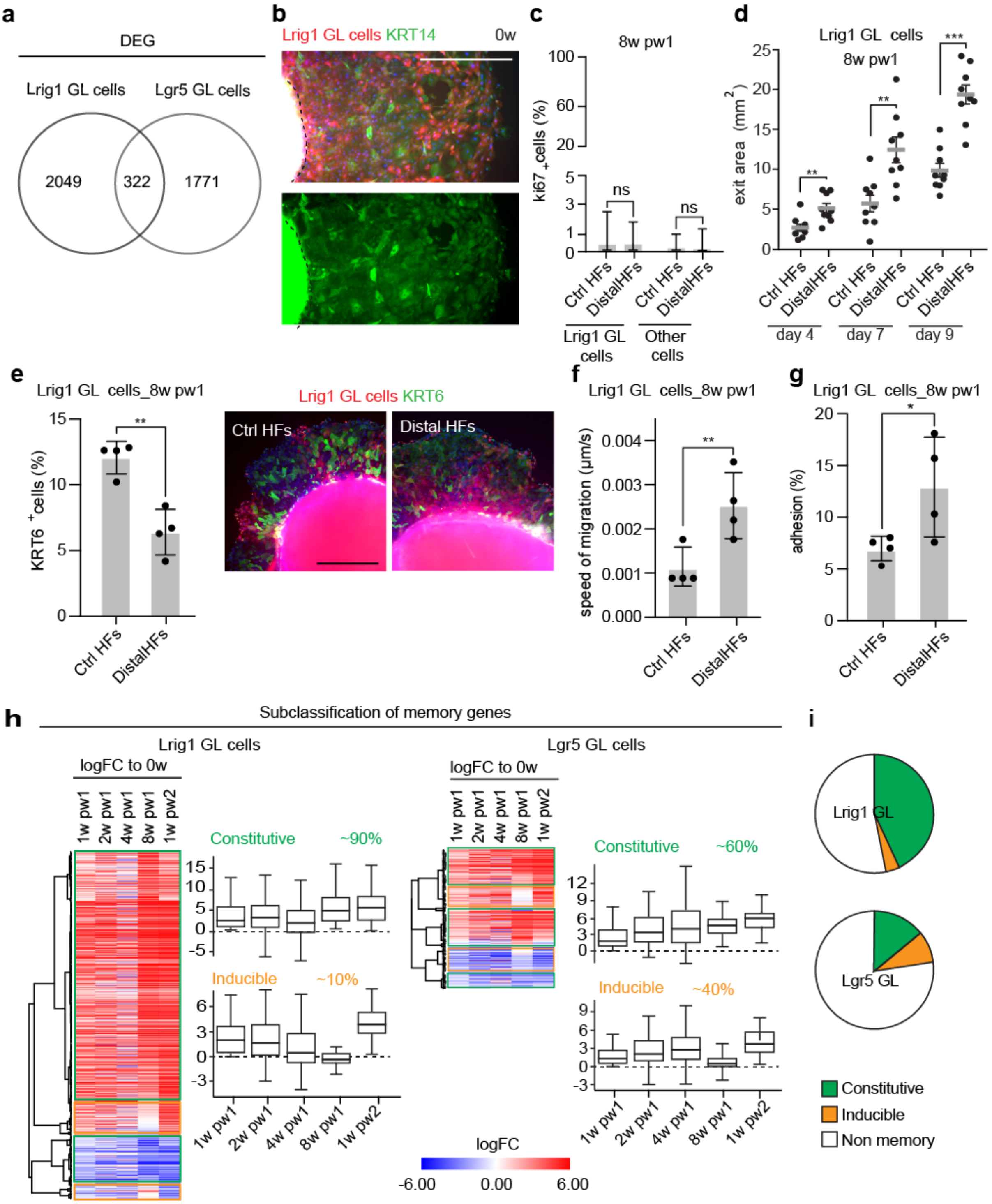

**Extended Data Fig.2: Transcriptome of Lrig1 GL cells suggests a new primed subpopulation arising after wound resolution. (a)** Venn diagram of unique and shared DEG between Lrig1 and Lgr5 GL cells. **(b, c)** Ex vivo culture of skin explant to assess migratory ability of epidermal cells. $tdTomato^+$ GL cells exiting from the skin explant are stained with KRT14 (green), marker for epidermal cells (b). Epidermal cells exiting skin explant do not proliferate. Percentage of $Ki67^+$ nuclei was measured 4 days after culture establishment. Mean with SEM is plotted. n = 9 explants (c). **(d)** Quantification of the exit area ($mm^2$) from the explant of Lrig1 GL. Mean with SEM is plotted. n=9 explants. **(e)** In cultured cells Krt6 positivity is inversely proportional to cell migration, consistently with the results from Wang et al. study[47]. Explants are stained after 4 days of culture with Krt6. Percentage of $Krt6^+$ in Lrig1 GL cells (left) and representative pictures are shown (right). n=4 explants. **(f)** Speed of migration (μm/s) in Lrig1 GL cells, as in Figure 1D, from time lapse migration assay. n=4 mice. **(g)** Percentage of cell adhesion of Lrig1 GL cells is calculated from the number of plated cells (day -1) and the counted cells at day 0. Lrig1 GL cells isolated from wound-educated distal HFs or Ctrl HFs are counted by flow cytometer ($tdTomato^+$). n=3 mice. **(h)** Heatmap of the memory genes showing as logFC of each time point respect to 0w. Memory genes can be divided into two subtypes, constitutive (green) and inducible (yellow), based on the expression at 8w pw1. **(i)** Pie chart illustrating the proportions of the memory genes subtypes.

P-value: *** 0.0001 to 0.001, ** 0.001 to 0.01, * 0.01 to 0.05, ns ≥ 0.05. Mean with SD is plotted if not differently indicated. Scale bars: 500 μm (b, e).

### Extended Data Fig.3

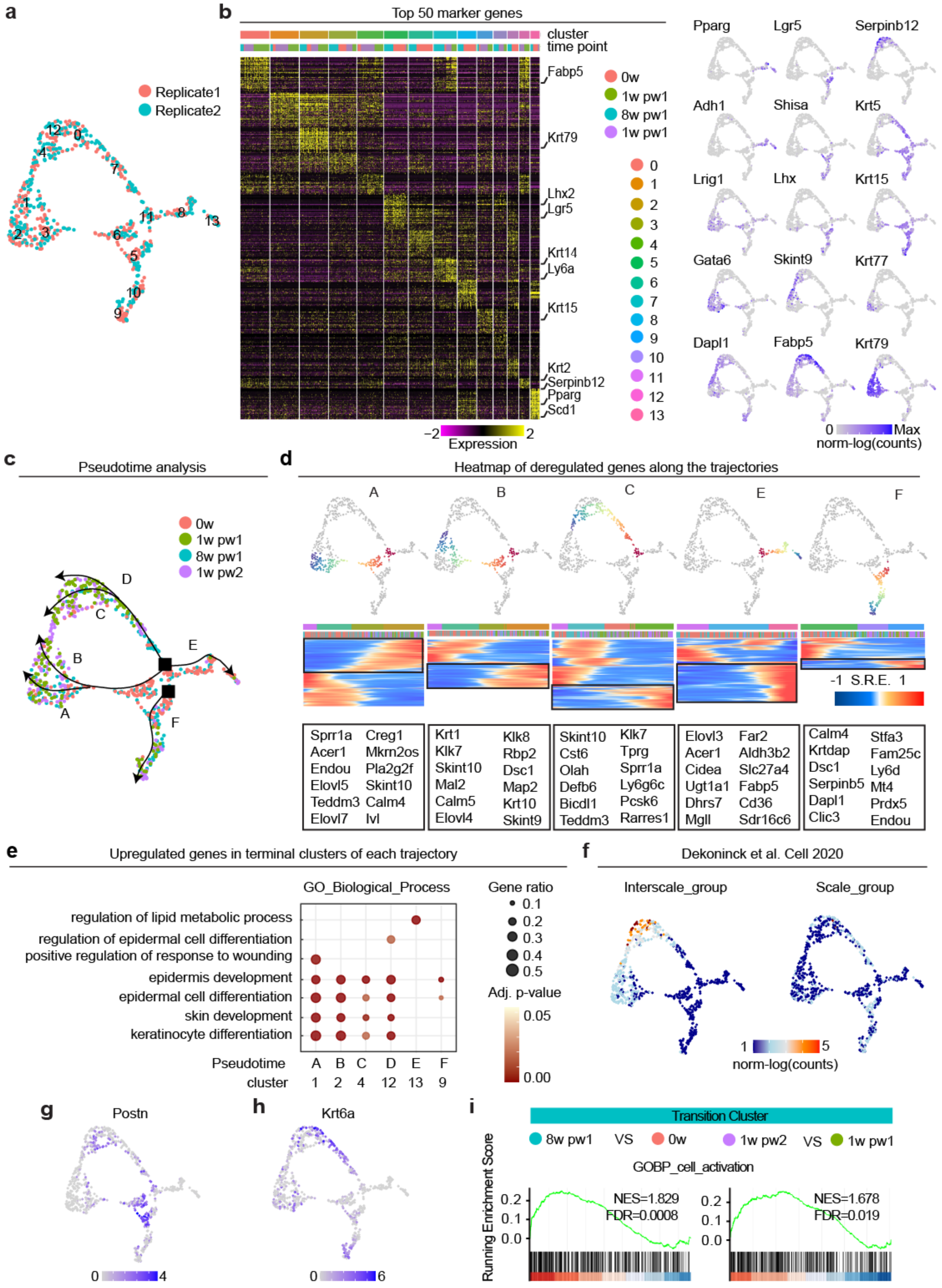

**Extended Data Fig.3: Single cell analysis assigned known epidermal niches to cell clusters. (a)** UMAP of scRNA-Seq replicates. To minimise biological divergences, around 100 cells per time point of each of the two replicates coming from a total of height wounded skin areas, were sequenced. **(b)** Heatmap representing the 50 top marker genes for each cluster (right) and UMAP of cells coloured by the expression of selected genes (left). Top bars indicate the clusters and the time points. **(c)** Trajectories identified by pseudotime analysis. The starting points (black square) are Cluster 11 (JZ) for the upper HF and Cluster 5 (bulge) for the lower HF. **(d)** Upper panel: plot of pseudotime trajectories and the associated heatmap with Smoothed Relative Expression (SRE). Lower panel: selected upregulated genes in the last cells of each trajectory are listed. Trajectory D is analysed alone in Figure 4. **(e)** GO enriched for induced genes in the terminal clusters of each trajectory. Gene ratio (number of genes in the pseudotime/ number of genes in the GO term) and adj. p-value are plotted. **(f)** UMAP of cells coloured by Interscale and Scale gene signature identified from Dekoninck et al. 2020. **(g, h)** Postn and Krt6a expression are plotted on the UMAP. Scale in log(counts). **(I)** GSEA ranking of the indicated gene signature using the differential gene expression estimates from the comparison 8w pw1 vs is 0w (left panel) and 1w pw2 vs 1w pw1 (right panel) in *Transition Cluster*.

### Extended Data Fig.4

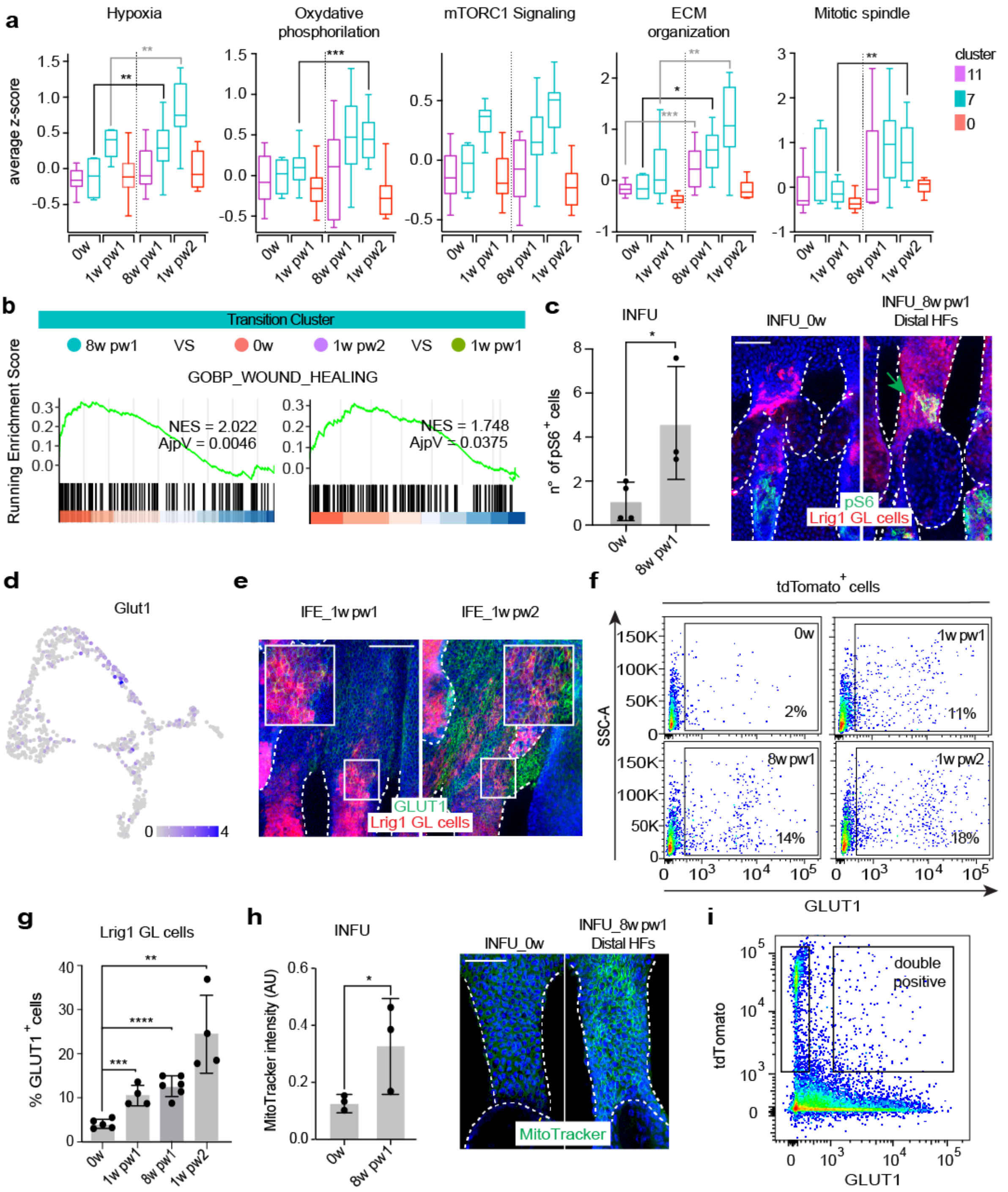

**Extended Data Fig.4: Characterisation of primed-memory cell state. (a)** Cells in pseudotime D are divided by time points and clusters and the average z-score of the indicated GO term is plotted in the whisker plot. Median with 25$^{th}$ and 75$^{th}$ percentiles is plotted. **(b)** GSEA ranking of the indicated gene signature using the differential gene expression estimates from the comparison 8w pw1 vs is 0w (left panel) and 1w pw2 vs 1w pw1 (right panel) in *Transition Cluster*. **(c)** Number of pS6 (Ser235-236) positive cells among Lrig1 GL cells in infundibulum (INFU) (left) and staining images at 0w or 8w pw1 (distal HFs ~ 5 mm from wound site) (right). n=3-4 wounds. **(d)** Glut1 expression levels are plotted on the UMAP. Scale in log(counts). **(e)** Whole-mount staining of Glut1 in 1w pw1 and 1w pw2. **(f, g)** Dot plot of Glut1$^{+}$ tdTomato$^{+}$ cells (f) and quantification (g) at the indicated time points. Gates are set on negative controls. n=4-6 wounds. **(h)** Quantification of MitoTracker index in INFU (left) and staining images (right) at 0w and 8w pw1 in distal HFs. n= 3 mice. **(i)** Gating strategy for tdTomato-Glut1 double positive cells sorting. P-value: **** < 0.0001, *** 0.0001 to 0.001, ** 0.001 to 0.01, * 0.01 to 0.05. Mean with SD is plotted if not differently indicated. Scale bars: 50 μm (c, e, h).

**Extended Data Fig.5**

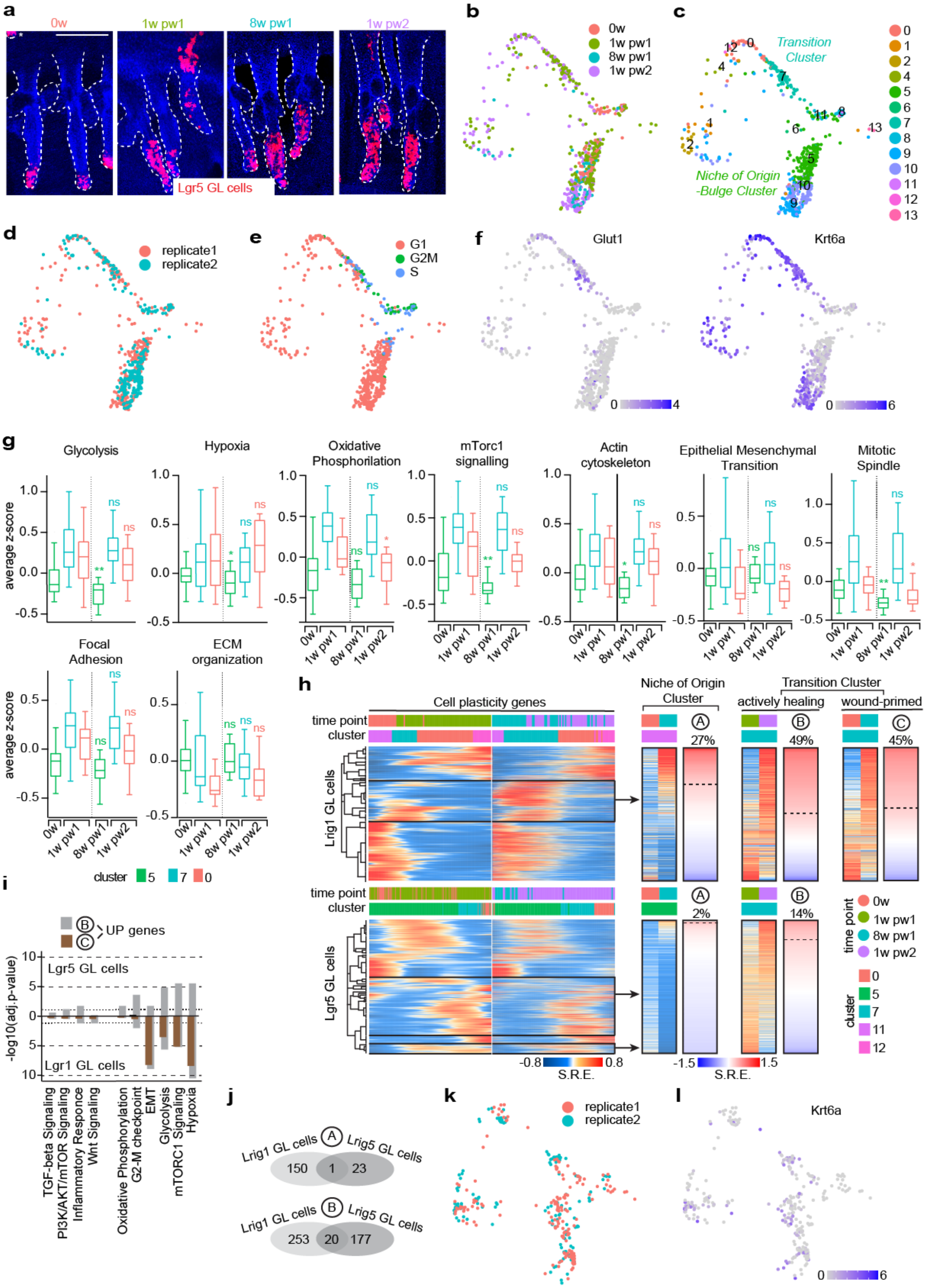

**Extended Data Fig.5: Comparison between single cell profiling of Lgr5 and Lrig1 lineages. (a)** Representative confocal pictures of Lgr5 GL HF triplets from tail skin at the indicated time points. Lgr5 GL cells exit from their niche, but they disappear by 8w pw1 if out of the bulge. Scale bar: 150 µm. **(b-f)** Single cell data from Lgr5 GL cells (0w, 1w pw1, 8w pw1 and 1w pw2). UMAP of cells coloured by time points (b), clusters (c), replicates (d) or cell-cycle phase (e) are shown. **(f)** Glut1 and Krt6a expression levels are plotted on the UMAP. Scale in log(counts). **(g)** Whisker plots of the average expression of each GO term enriched in Lrig1 GL single cells. No significant enrichment is present. Median with $25^{th}$ and $75^{th}$ percentiles is plotted. **(h)** Left panel: Smoothed Relative Expression (SRE) of deregulated genes in the clusters 0, 7, 11 and 12 for Lrig1 GL cells (up) and 5, 7 and 0 for Lgr5 GL cells (down). Clusters and time points are indicated above, and transiently induced cell plasticity genes are highlighted (black rectangle), as in Figure 4B. Right panel: the pattern and the percentage of cell plasticity genes are shown for: “A” cells in the *Niche of Origin Cluster* (cluster 5 for Lgr5 GL and cluster 11 for Lrig1 GL cells); “B” actively healing cells at 1w pw1 and 1w pw2 in *Transition Cluster*, “C” wound-primed cells at 8w pw1 in *Transition Cluster* only exists in Lrig1 GL cells. **(i)** GO analysis for the deregulated genes in cell subsets “B” and “C” in Lrig1 and Lgr5 GL cells. -log10 of the adj. p-value is reported. Dashed lines represent significance. **(j)** Venn diagram of the genes in A and B between Lgr5 and Lrig1 GL populations. **(k)** UMAP of cells coloured by replicates. **(l)** Krt6a expression levels. Scale in log(counts).

**Extended Data Fig.6**

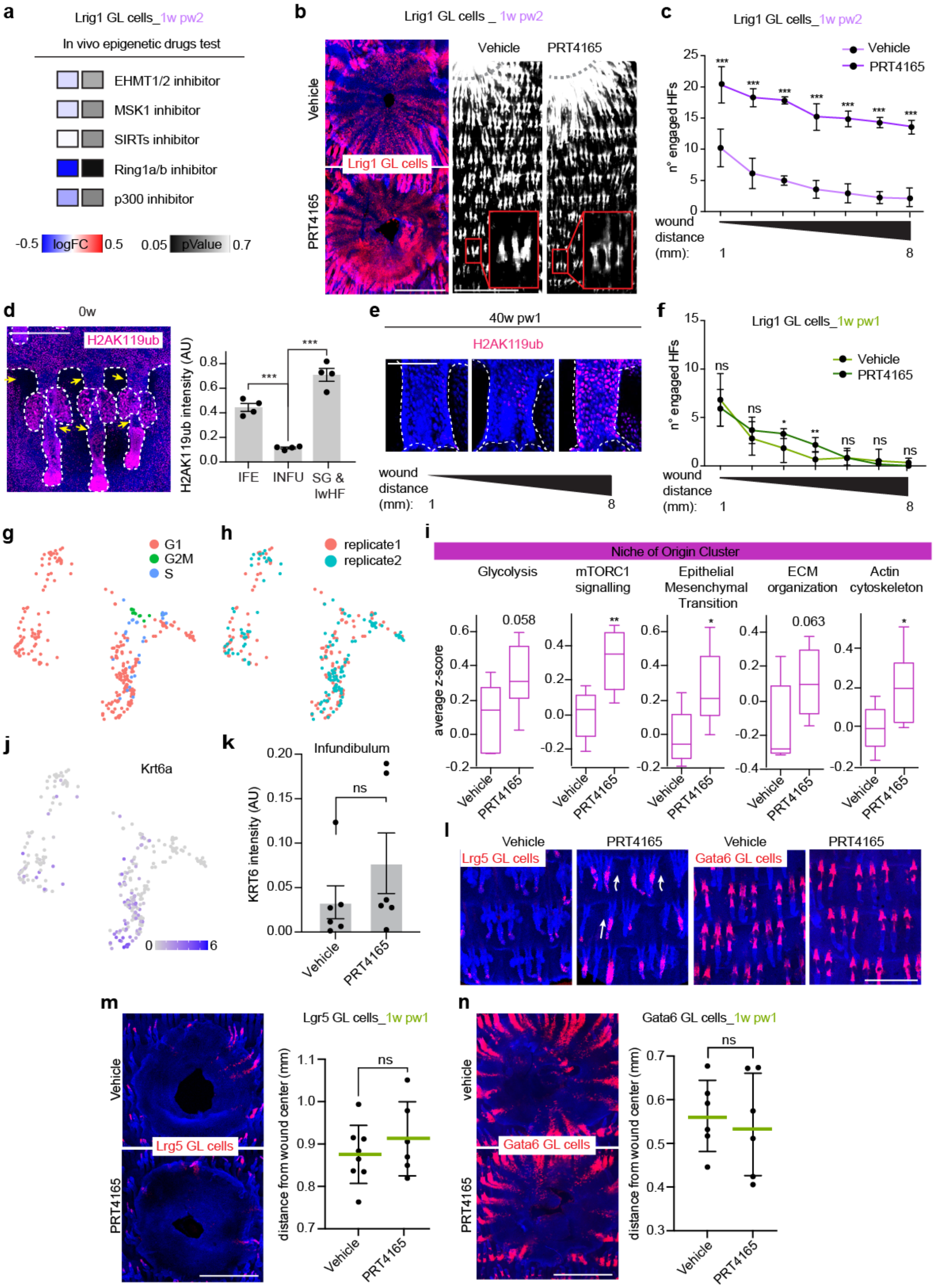

**Extended Data Fig.6: Epigenetic characterisation and manipulation of wound-primed memory. (a)** Summary of drug screening results at 1w pw2. 8w pw1 mice are treated 3 times with the reported inhibitors before wound induction. 1w pw2 distance from wound centre is quantified and plotted as logFC to vehicle treated mice. Only the Ring1a/b inhibitor (PRT4165) is significantly able to increase wound closure rate. n=4-9 mice. **(b)** Representative pictures of the effect of PRT4165 on wound closure (left) and distal HFs engagement in Lrig1 GL tdTomato$^{+}$ mice at 1w pw2. **(c)** Number of engaged HFs at 1w w2 in different distances from wound after vehicle/ PRT4165 treatment. n=7-8 wounds. **(d)** Homeostatic distribution of H2AK119ub in HFs and relative quantification in interfollicular epidermis (IFE), infundibulum (INFU) and sebaceous gland (SG) and lower HF (lwHF). INFU is the compartment with lower levels of H2AK119ub in homeostasis. Scale bar 150µm. n=4 mice. **(e)** H2AK119ub index in 40 weeks post wound (40w pw1) homeostasis at different distances from wound. Scale bar: 50 µm. **(f)** Number of engaged HFs at 1w pw2 in different locations, up to 8 mm away from wound after vehicle/PRT4165 treatment. n=6 wounds. **(g, h)** Single cell data from 0w Lrig1 GL cells treated with vehicle/ PRT41645. UMAP of cells coloured by cell-cycle phases (g) or cluster (h). **(i)** Whisker plots of the average expression for each GO term enriched in Lrig1 GL single cells. Median with 25$^{th}$ and 75$^{th}$ percentiles is plotted. **(j)** Expression of Krt6a is plotted on the UMAP. Scale in log(counts). **(k)** Quantification of Krt6 staining in the infundibulum in vehicle/PRT4165 treated epidermis. n=6 wounds. **(l)** Confocal pictures of Lgr5 or Gata6 GL cells showing the macroscopic effect of PRT4165 treatment. **(**m, n) Epidermal whole-mount showing GL tdTomato+ cells at wound site (left) and quantification of distance from the wound centre (right) in vehicle vs PRT4165 treated samples at 1w pw1 in Lgr5 (m) or Gata6 GL epidermis (n). n=6-8 wounds. P-value: *** 0.0001 to 0.001, ** 0.001 to 0.01, * 0.01 to 0.05, ns ≥ 0.05. Mean with SD is plotted if not differently indicated. Scale bars: 1mm (b, g, h, i).

**Extended Data Fig.7**

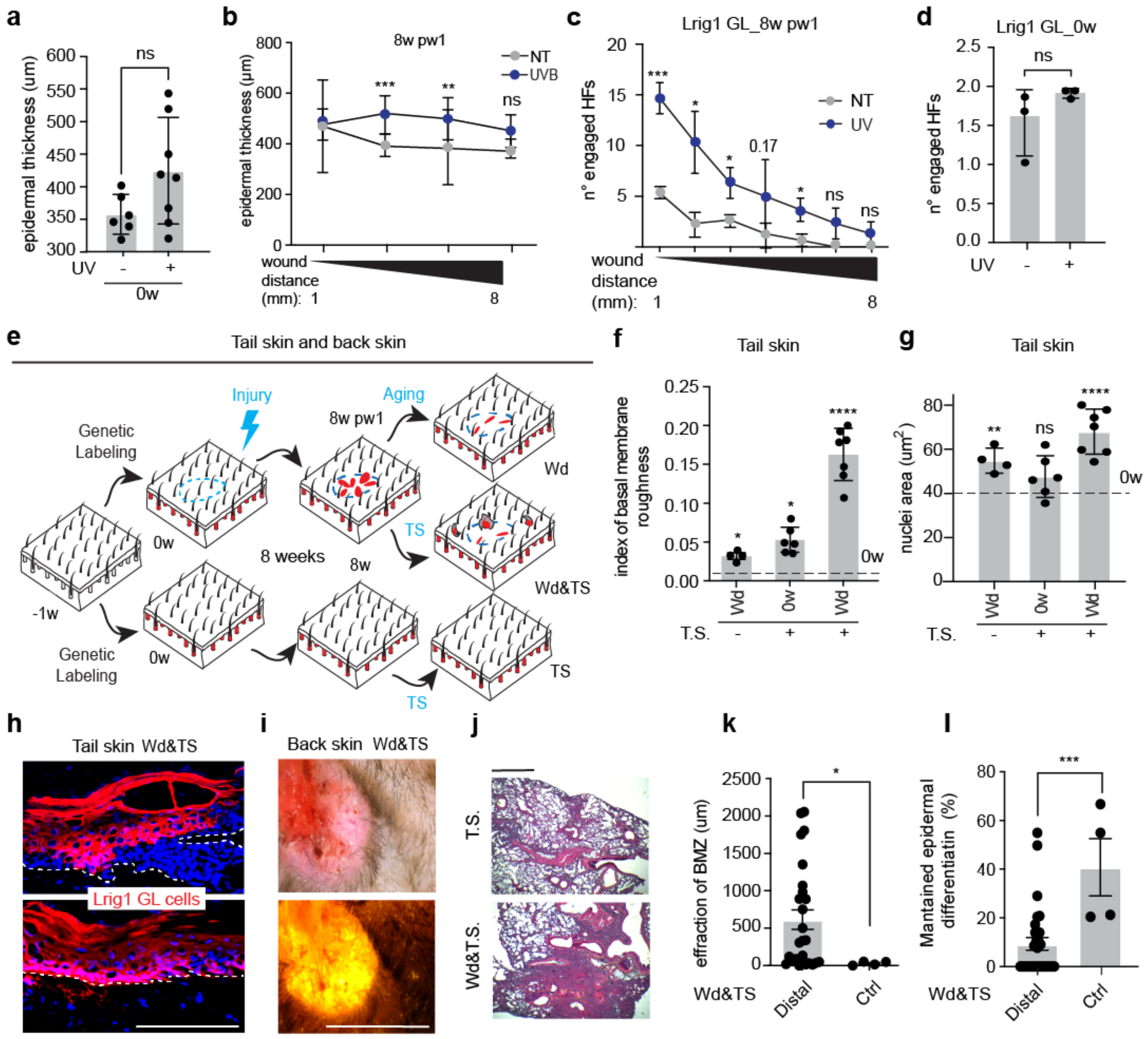

**Extended Data Fig.7: Healed skin is more sensitive to tumour onset. (a, b)** Effect of acute UV irradiation on epidermal thickening at 0w (n=6-8 mice) (a) or 8w pw1 (n =4-11 mice) (b), where different distances from wound site were considered. NT= not treated. **(c, d)** Effect of acute UV irradiation on the number of engaged HF at 8w pw1, at different wound distances (c) or 0w (d). n=3. **(e)** Scheme of tumorigenesis protocol in Lrig1 GL mice: (-1w) genetic labelling; (0w) injury or not; (8w) 8w weeks after, tumorigenic stimuli (TS) started (tail or back skin). Samples are wounded only (Wd), wounded with TS (Wd&TS) or only treated with TS (TS). **(f, g)** Characterisation of early SCC phenotype in tail skin. Quantification of membrane roughness in Wd (n=4), TS (n=6) and Wd&TS (n=7) mice (f) and nuclei area is quantified in the same sample (g). **(h)** Representative pictures of Lrig1 GL cells in early SCC (eSCC) lesions. **(i)** Macroscopic picture of back skin SCC in Lrig1 GL tdTomato$^+$ mice in brightfield (left) and red channel (right). Scale bar: 1mm. **(j)** H&E staining of lung metastasis derived from back skin primary tumours. **(k, l)** Characterisation of tumours developed in Distal or Ctrl zones as described in Figure 7. Effraction of the basal membrane zone (BMZ) informative for tumour invasiveness. Statistics: Mann-Whitney t-test. n=4 Ctrl, n=27 Distal tumours (k); percentage of maintained epidermal differentiation was evaluated over the total tumour area. n = 4 Ctrl, n = 30 Distal tumours (l).

P-value: **** < 0.0001, *** 0.0001 to 0.001, ** 0.001 to 0.01, * 0.01 to 0.05, ns ≥ 0.05. Mean with SD is plotted if not differently indicated. Scale bars: 300 μm (h, j).
